# Allosteric Regulation of Human Plastins

**DOI:** 10.1101/2021.12.01.470822

**Authors:** Christopher L. Schwebach, Elena Kudryashova, Richa Agrawal, Weili Zheng, Edward H. Egelman, Dmitri S. Kudryashov

## Abstract

Plastins/fimbrins are conserved actin-bundling proteins contributing to motility, cytokinesis, and other cellular processes by organizing actin assemblies of strikingly different geometries as in aligned bundles and branched networks. We propose that this unique ability stems from an allosteric communication between plastins’ two actin-binding domains (ABD1/2) engaged in a tight spatial association. We found that although ABD1 binds actin first, ABD2 can bind to actin three orders of magnitude stronger if not inhibited by an equally strong allosteric engagement with ABD1. Binding of ABD1 to actin lessened the inhibition, enabling weak bundling within aligned bundles. A mutation mimicking physiologically relevant phosphorylation at the ABD1-ABD2 interface strongly reduced their association, dramatically potentiating actin cross-linking. Cryo-EM reconstruction revealed the ABD1-actin interface and enabled modeling of the plastin bridge to confirm domain separation in parallel bundles. The characteristic domain organization with a strong allosteric inhibition imposed by ABD1 on ABD2 allows plastins to tune cross-linking, contributing to the assembly and remodeling of actin assemblies with different morphological and functional properties defining the unique place of plastins in actin organization.

## INTRODUCTION

Plastins, also known as fimbrins, are a family of actin-bundling proteins conserved throughout eukaryotic life and involved in cell migration, cytokinesis, and endocytosis ^1, 2^. Vertebrates express three tissue-specific plastin isoforms. Plastin 1 (PLS1; a.k.a. I-plastin) is primarily found in microvilli in the epithelial brush border of the intestine and kidneys ^3, 4^, and in the microvilli-like stereocilia of the inner ear hair cells ^5, 6^. Genetic defects in PLS1 gene are linked to hearing loss ^7, 8^. Plastin 2 (PLS2; a.k.a. L-plastin, LCP1, LPL) is expressed in hematopoietic cells contributing to immune cell activation, migration, and invasion ^9–15^. Congenital diseases associated with PLS2 mutations are not known, but, intriguingly, PLS2 is ectopically expressed in ∼70% of epithelial cancers, where it localizes to the leading edge and invadopodia contributing to metastatic capabilities of these cells ^16–20^. Plastin 3 (PLS3; a.k.a. T-plastin) is the most ubiquitous of the three human isoforms expressed by most solid tissues, where it is enriched at the cell edge, specialized adhesion contacts ^21^, distal parts of filopodia ^22^, sites of endocytosis ^23^, and in the contractile actin cortex formed upon membrane blebbing repair ^24^. Mutations in PLS3 lead to severe X-linked hereditary osteoporosis with bone fragility ^25–35^. Congenital osteoporosis develops upon deletion and truncations in PLS3 gene (reviewed in ^36^), but also due to point mutations resulting in PLS3 variants, which either lose the ability to bundle F-actin or display perturbed sensitivity to Ca^2+ 37^. By contributing to endocytosis ^38^ and migration, PLS3 plays neuroprotective roles by lessening the toxicity of pathogenic poly-glutamine-containing proteins ^39^, alleviating symptoms of spinal muscular atrophy ^40, 41^, and coordinating neuronal cell migration in embryogenesis ^42^. Furthermore, PLS3 is also recognized as an oncogene, whose ectopic expression in hematopoietic cells ^43, 44^ and overexpression in cells of solid tissues promote cancer cell proliferation, motility, survival, and drug resistance ^45, 46^.

Plastins belong to a superfamily of calponin-homology (CH)-domain-containing actin cytoskeleton organizers, which also include filamins, spectrins, α-actinins, IQGAP, and other actin-binding proteins ^47^. Most members of the superfamily contain only one actin-binding domain (ABD) and must form dimers or higher-order oligomers to tether actin filaments into bundles and meshwork. Plastins are unique in having two ABDs within the same polypeptide chain (Fig. 1a). Each ABD consists of two CH-domains. In addition, all plastins contain an N-terminal regulatory domain (RD) comprised of two EF-hands followed by an EF-hand-binding motif [EBM; ^48, 49^] and a long, disordered linker connecting the regulatory domain to the ABD1-ABD2 core (Fig. 1a). In mammalian plastins, the actin-bundling activity is inhibited upon Ca^2+^ binding to EF-hands ^4, 40, 50^, albeit some non-mammalian fimbrins are insensitive to Ca^2+ 51–53^.

**Figure 1.**
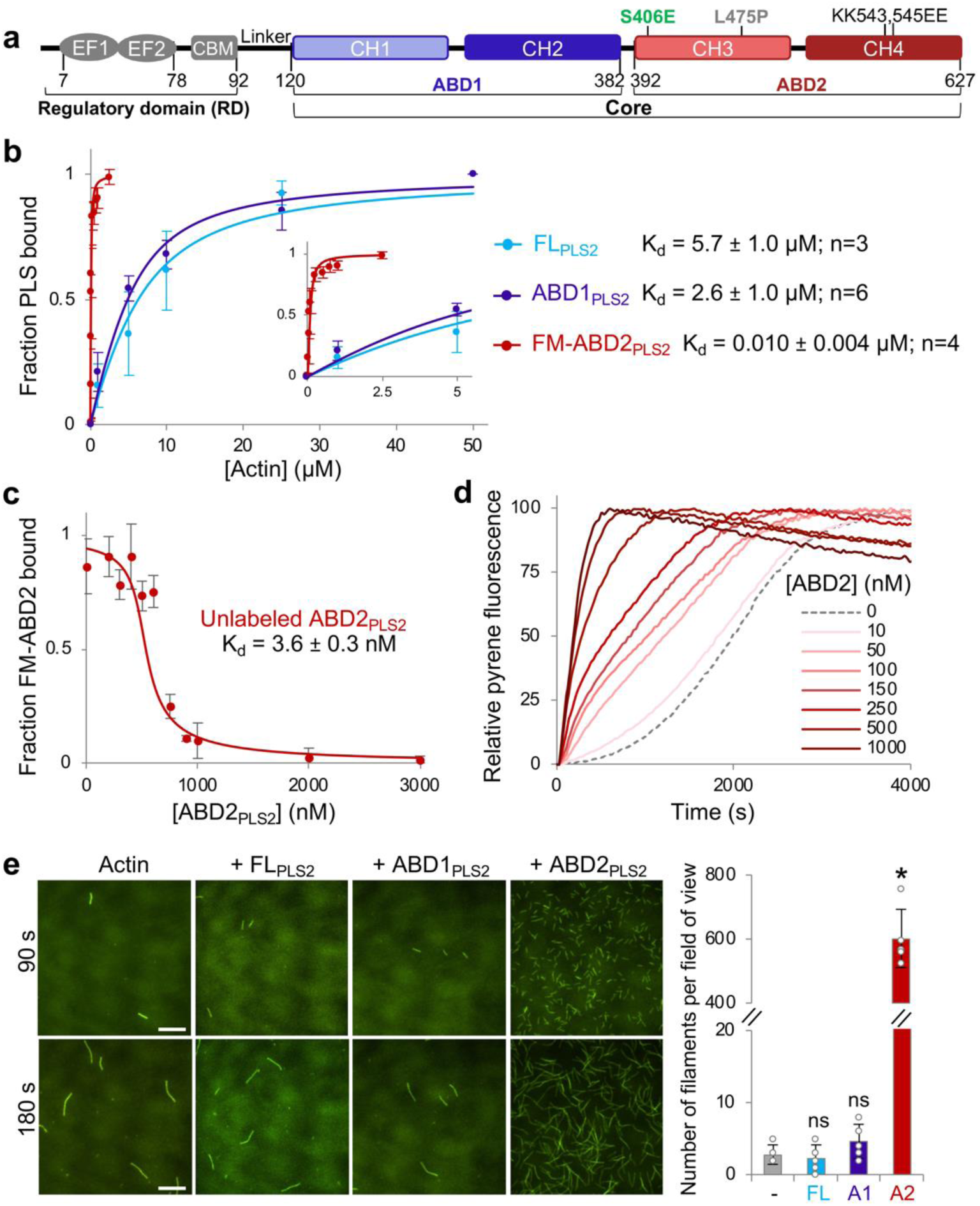
ABD2 of PLS2 binds to F-actin with nanomolar affinity and nucleates actin. **(a)** A schematic representation of plastin domain organization. Regulatory domain (RD) consisting of two EF-hand motifs (EF1 and 2) is connected through a flexible linker to the PLS2 core comprising two actin-binding domains (ABD1 and 2), each consisting of two calponin-homology motifs (CH1-4). Numbers represent amino acid numbering in the PLS2 sequence. Relevant mutations are indicated. **(b)** F-actin binding affinities of the full-length PLS2 (FL_PLS2_) and ABD1_PLS2_ were assessed by F-actin co-sedimentation assays; F-actin binding affinity of ABD2_PLS2_ was measured by FA. The inset zooms in the data range 0 – 5 µM of actin. Error bars represent SD of the mean; number of repetitions for each group (n) is indicated in the figure. **(c)** F-actin binding affinity of unlabeled ABD2_PLS2_ was assessed by FA competition assays. Error bars represent the SD of the mean; n=3. **(d)** Bulk pyrene-actin polymerization was monitored in the absence or presence of increasing concentrations of ABD2; representative normalized fluorescence traces are shown. **(e)** Actin nucleation activities of FL_PLS2_ and its individual ABDs (50 nM final concentration) were tested by TIRFM. Representative micrographs at 90 and 180 s after the initiation of actin polymerization are shown; scale bars are 10 µm. The graph shows the number of actin filaments present in a field of view at 90-s time point. Error bars represent the SD of the mean; n=5. ANOVA followed by multiple comparison tests with Bonferroni correction was applied: asterisk indicates statistically significant difference (*p<0.0167); individual *p*-values are 0.58 (FL_PLS2_ (FL)), 0.18 (ABD1_PLS2_ (A1)), and *4.6×10^-7^ (ABD2_PLS2_ (A2)), each compared to the actin control (-).

The current understanding of the plastin organization is based on crystal structures of the ABD1-ABD2 cores of yeast and plant fimbrins ^30^. For vertebrate plastins, only structures of individual ABD1 ^54^ and CH4 of ABD2 (PDB:1WJO, 2D85) were resolved by X-ray crystallography and solution NMR, respectively, and the interaction of ABD2 (but not ABD1) with actin was revealed by cryo-electron microscopy (cryo-EM) reconstructions ^37, 55^. EM of negatively stained two-dimensional actin arrays assembled on lipid surfaces ^56^ revealed the overall architecture, spacing, and polarity of plastin-tethered bundles. It was found that actin filaments in the arrays are assembled in tight parallel bindles with 120-Å spacing between centers of neighboring filaments. Unfortunately, the resolution of the reconstruction (∼37 Å) did not allow for unambiguously defining the domain orientation and the interaction mode of ABDs with actin. Structures of the EF-hand domain with and without Ca^2+^ were determined by solution NMR ^49^, but the RD location relative to the ABD1-ABD2 core has not been revealed by high-resolution structural methods. Analysis of PLS3 pathogenic variants linked to hereditary osteoporosis revealed their aberrant sensitivity to Ca^2+^ and pointed to the localization of RD at the interface between ABD1 and ABD2, in the region enriched by several disordered loops ^37^. Accordingly, the mechanism of plastins’ regulation was proposed: while binding of ABD1 to actin remains unaffected by Ca^2+ 48^, the RD in the Ca^2+^-bound state locks ABD2 in the inhibited conformation preventing the transition of the CH4 subdomain into a position where it does not clash with actin. In the proposed model, it remained unclear whether the inhibitory effect on ABD2 is imposed strictly by the RD or the inhibition originates from ABD1 and only is regulated by the RD.

Here, we report an overarching mechanism of plastin regulation that is mediated by a strong allosteric inhibition imposed by ABD1 on ABD2. We employed covalent cross-linking to stabilize an otherwise insufficiently stable complex of ABD1 with F-actin and obtained F-actin filaments decorated by ABD1. Cryo-EM reconstruction revealed that binding of ABD1 to actin roughly recapitulates that of ABD2, although with a smaller footprint area of CH1 on actin compared to CH3 of ABD2, and CH2 separated from actin further away than CH4 of ABD2. Superposition of the actin-bound ABD1 structure with that of unbound ABD1-ABD2 core confirmed that moderate reorientation of ABD2 is needed to avoid clashes with actin. Accordingly, we found that the ABD2 inhibition by ABD1 is slightly attenuated upon binding of ABD1 to F-actin, allowing engagement of ABD2 in weak actin bundling. The extent of the inhibitory allosteric engagement between ABD1 and ABD2 can be further weakened by a mutational mimicking of physiologically relevant phosphorylation of the Ser406 residue at the ABD1-ABD2 interface. As a result, the reduced allosteric engagement between ABDs strongly increased the bundling potential of the protein. Ca^2+^ did not affect the interaction between ABD1 and ABD2, suggesting that it regulates actin bundling independently of the allosteric mechanism. Accordingly, Ca^2+^ retained the ability to attenuate actin bundling by the S406E plastin mutant, but to a lesser degree than bundling by WT protein. We propose that the discovered allosteric inhibition mechanism enables regulating the strength of actin cross-linking by plastins in a broad range contributing to the assembly and remodeling of various high-order actin structures with different morphological and functional properties. We suggest that this ability is a key feature of plastins that distinguishes them from other CH-domain actin organizers and defines their unique place in the organization of the actin cytoskeleton.

## RESULTS

### ABD2 binds to F-actin orders of magnitude stronger than the full-length protein

ABD1 of human PLS3 (ABD1_PLS3_) binds F-actin with a K_d_ in a low-micromolar range, similar to that of full-length (FL) PLS3 (FL_PLS3_) ^48^. Similarly, co-sedimentation assays revealed that ABD1_PLS2_ binds to F-actin only slightly better than FL_PLS2_ (K_d_ of 2.6 ± 1.0 µM and 5.7 ± 1.0 µM, respectively; Fig. 1b). To evaluate the contribution of both domains to actin binding, we attempted to produce recombinant ABD2 of the human plastin isoforms. Of all three, only ABD2_PLS2_ was soluble, stable, and existed in solution as a monomeric protein as confirmed by gel filtration chromatography and analytical sedimentation (Supplementary Fig. 1a-c). In contrast, ABD2 from PLS1 and PLS3 could not be purified due to their intrinsically low stability either alone or as a fusion with a folding-promoting maltose-binding protein ^57^. To our surprise, isolated ABD2_PLS2_ bound F-actin much stronger than ABD1_PLS2_ or the FL_PLS2_. Since the affinity was too high to be accurately assessed by F-actin co-sedimentation assays, fluorescence anisotropy (FA) experiments were employed to reveal that fluorescein-labeled ABD2_PLS2_ (FM-ABD2) binds F-actin with a K_d_ in the nanomolar range (9.9 ± 3.7 nM; Fig. 1b), while unlabeled ABD2_PLS2_ binds even tighter (K_d_ = 3.6 ± 0.3 nM; Fig. 1c). Therefore, the affinity of isolated ABD2 to actin is ∼1,500 times stronger than that of full-length PLS2.

Given that ABD2 binds F-actin at the interface between two long-pitch actin subunits ^37^, we hypothesized that such a strong binding might stabilize the longitudinal dimer and promote nucleation. Indeed, in pyrenyl-actin polymerization assays, low-nanomolar concentrations of ABD2_PLS2_ potently increased bulk actin polymerization rate (Fig. 1d). A drastically increased number of filaments in the presence of ABD2_PLS2_, but neither ABD1_PLS2_ nor FL_PLS2_, observed by total internal reflection fluorescence microscopy (TIRFM; Fig. 1e) confirmed that faster polymerization is indeed due to actin nucleation.

Polymerization of actins with perturbed filament-forming interfaces can be rescued by filament interface-stabilizing factors ^58, 59^. Intermolecular covalent cross-linking of actin monomers by actin cross-linking domain (ACD) toxin brings actin subunits into oligomers that do not polymerize as their longitudinal contacts are disturbed by artificial covalent bonds ^60^. The polymerization of ACD-cross-linked actin oligomers can be rescued, however, if the perturbed contacts are compensated by enforcing the remaining contacts by phalloidin or reshaping the contacts by cofilin ^60, 61^. Accordingly, in agreement with its ability to staple adjacent subunits ^37^ by the strong ABD2-actin contacts, ABD2_PLS2_ effectively rescued polymerization of ACD-cross-linked actin oligomers (Supplementary Fig. 1d). The above results demonstrate that ABD2 in isolation can bind to F-actin with very high affinity and potently nucleate actin, *i.e.*, possesses abilities suppressed in the full-length protein context.

### ABD1 and ABD2 of plastins interact *in trans* with nanomolar K_d_

The potent nucleating ability of ABD2_PLS2_ was not reproduced by full-length plastin (Fig. 1e), suggesting the suppression of the nucleation capacity of ABD2 in the full-length protein context, potentially due to its interaction with ABD1. Accordingly, added *in trans*, ABD1_PLS2_ inhibited the nucleating ability of ABD2_PLS2_ in a dose-dependent manner (Fig. 2a). In FA experiments, ABD1_PLS2_ bound to ABD2_PLS2_ with a high affinity (K_d_ = 11.4 ± 0.7 nM in FA competition assays; Fig. 2b), whereas the ABD1 construct with the regulatory domain (RD-ABD1_PLS2_) bound ABD2_PLS2_ even stronger, irrespective of the presence of Ca^2+^ (K_d_ = 3.0 ± 0.9 nM in EGTA vs K_d_ = 2.9 ± 0.5 nM in Ca^2+^; Fig. 2c). Therefore, RD contributes to the interaction between the actin-binding domains, in agreement with its recent mapping to the loop-rich region at the ABD1-ABD2 interface ^37^. The ABD1-ABD2 affinity was nearly identical in 30 and 130 mM KCl (Supplementary Fig. 2), suggesting a dominant role of hydrophobic surfaces in this interaction.

**Figure 2.**
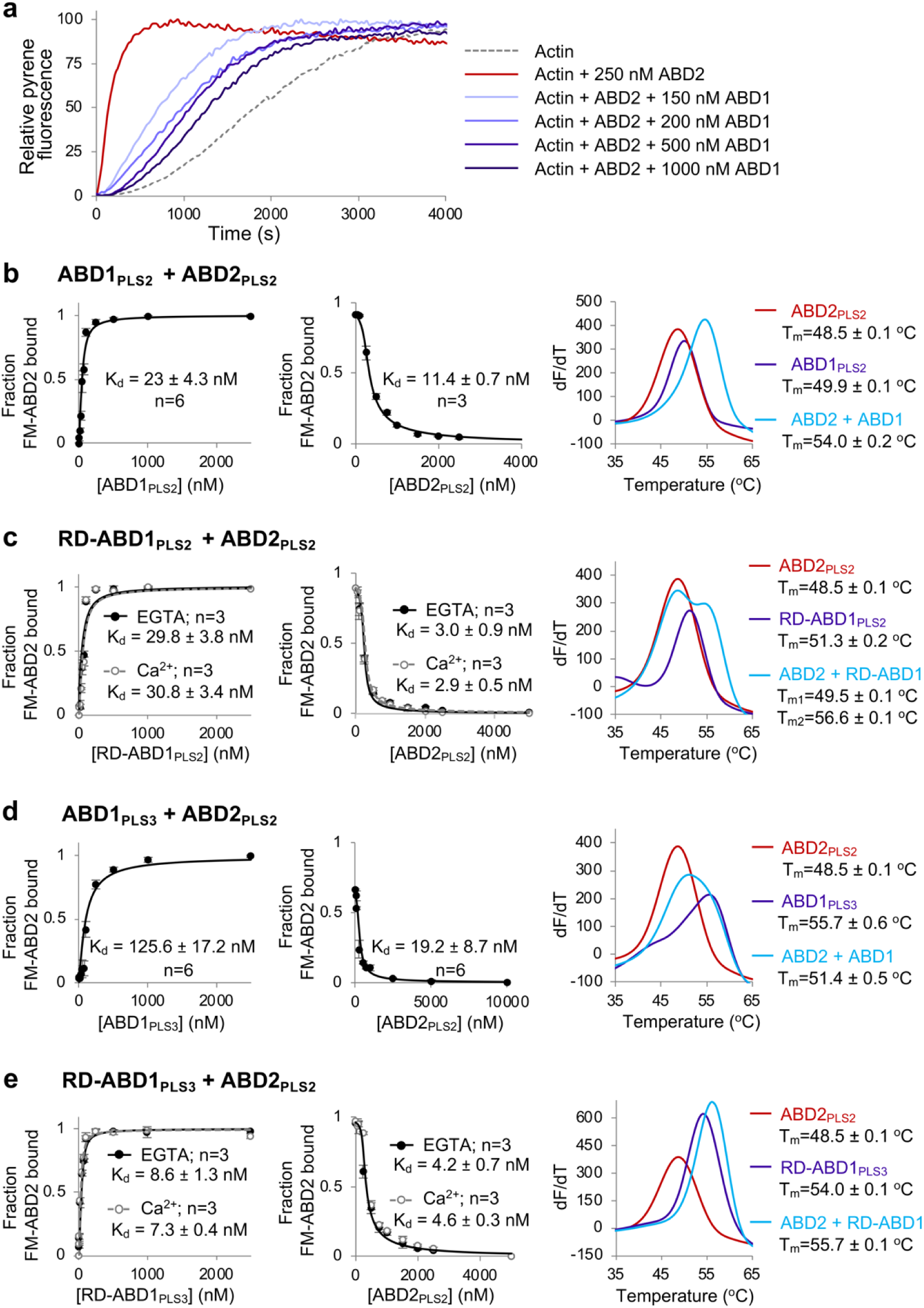
ABD1 of PLS2 interacts with and suppresses ABD2. **(a)** Effects of increasing concentrations of ABD1_PLS2_ on the ABD2_PLS2_-mediated acceleration of actin polymerization was monitored by bulk pyrene-actin polymerization assays; representative normalized fluorescence traces are shown. **(b-e)** Binding affinities of the individual ABDs of PLS2 and PLS3 to each other *in trans* were measured by fluorescence anisotropy using FM-ABD2_PLS2_ (left panels) and confirmed in FA competition assays using unlabeled ABD2_PLS2_ (middle panels). Error bars represent the SD of the mean; number of repetitions (n) is indicated in the figures. Melting temperatures (T_m_s) of the individual ABDs and upon mixing with each other were tested by DSF (right panels): the graphs show the first derivatives (-dF/dT) of melting curves (average of n=3).

Increase in thermostability often accompanies formation of protein complexes and can be detected by differential scanning fluorimetry (DSF) ^62^. Accordingly, the *in trans* ABD1-ABD2 complexes were more stable than individual domains (Fig. 2b,c), not however reaching the stability of FL_PLS2_ (T_m_ = 65 °C; Figs. 3 and 5b).

**Figure 3.**
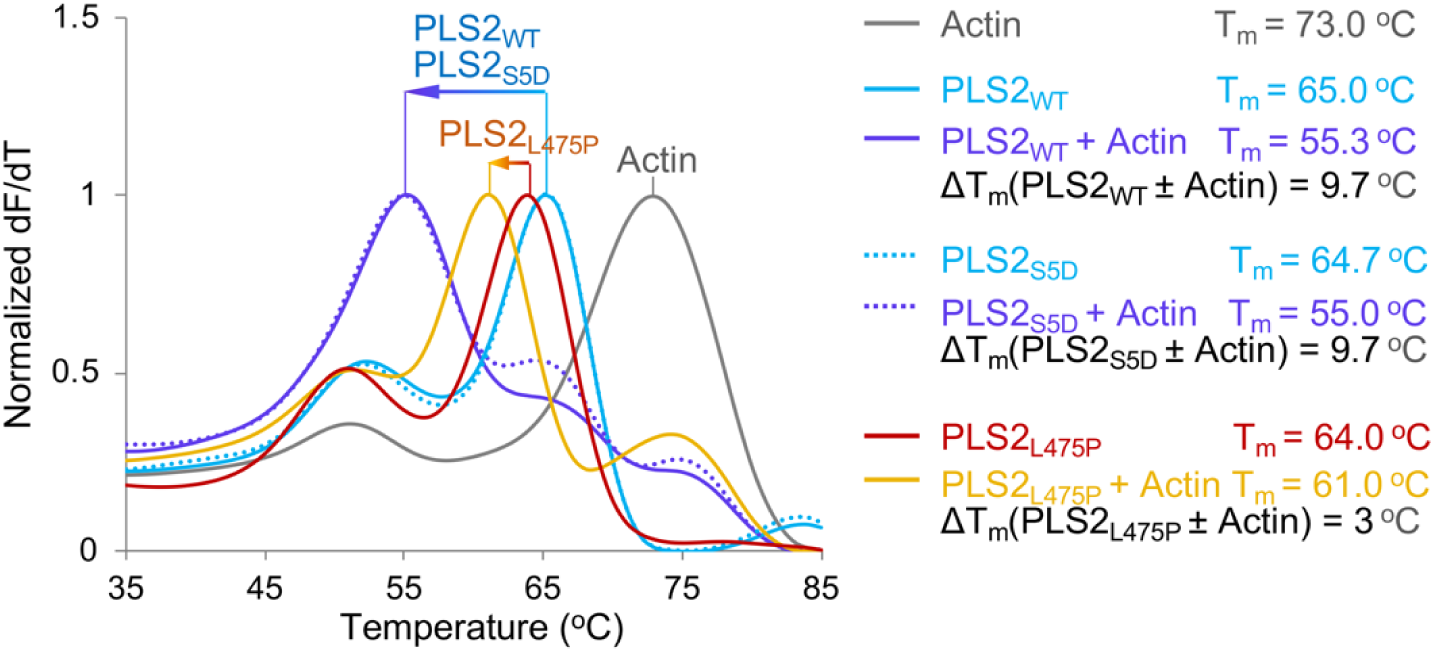
F-actin binding releases the ABD1-imposed suppression of ABD2. Melting temperatures (T_m_s) of bundling-competent (WT and S5D) and bundling-incompetent (L475P) full-length PLS2 constructs were assessed by DSF in the absence or presence of F-actin. The normalized first derivative melting curves (-dF/dT) represent the average data of n=3. The arrows indicate the direction/magnitude of the shifts of PLS2 melting peaks upon F-actin binding.

Due to reduced stability and prompt degradation, neither ABD2_PLS1_ nor ABD2_PLS3_ could be characterized. Yet, ABD1_PLS3_ and RD-ABD1_PLS3_ bound to ABD2_PLS2_ with high affinities similar to those of ABD1_PLS2_ and RD-ABD1_PLS2_, respectively (Fig. 2d,e), suggesting that the RD-ABD1/ABD2 interface is highly preserved in plastin isoforms.

Therefore, ABD1 and ABD2 of plastins interact with each other with high affinity comparable to that of isolated ABD2 to F-actin, suggesting that ABD1/ABD2 interaction suppresses the ABD2 high-affinity binding to F-actin.

### F-actin binding weakens ABD1-imposed inhibition of ABD2

A strong inhibitory influence of ABD1 on ABD2 reconciles the apparent paradox that the domain with a much higher affinity to actin (*i.e.,* ABD2) binds actin only after the lower-affinity ABD1 domain is bound ^48^. Given that, even *in trans*, the strength of the inhibitory ABD1/ABD2 interaction is nearly matching that of the bundling-conferring ABD2/F-actin interaction (Figs. 2b,c and 1b,c), the inhibitory component must be prevailing when ABD1 and ABD2 are parts of the same protein. We hypothesized, therefore, that for bundling to happen, the inhibition must be weakened by a mechanism universally applicable to all plastins. Since binding of ABD1 to actin precedes that of ABD2 ^48^, we tested whether the interaction of ABD1 with actin may contribute to the inhibition release. Since separated ABD1 and ABD2 (T_m_ = 48.5 - 51.3 °C; Fig. 2b,c) are less stable than the FL_PLS2_ (T_m_ = 65 °C; Figs. 3 and 5b), we should expect destabilization of PLS2 upon binding to F-actin due to the anticipated ABD1-ABD2 disentanglement.

To distinguish the effects of the domain separation caused by binding via ABD1 from those resulting from binding via both domains, we reproduced the osteoporosis L478P_PLS3_ mutation that blocks ABD2 binding to actin ^37^ in PLS2 (L475P_PLS2_). We confirmed that, similarly to L478P_PLS3_, L475P_PLS2_ was able to bind (presumably through intact ABD1) but unable to bundle F-actin effectively regardless of the presence of Ca^2+^ (Supplementary Fig. 3). In agreement with our hypothesis, the T_m_ of L475P_PLS2_ decreased by ∼3 °C upon binding to F-actin (Fig. 3), suggesting a modest disengagement of the domains. In contrast, wild-type (WT) PLS2 (PLS2_WT_) and S5D PLS2 mutant (PLS2_S5D_) mimicking the Ser5 phosphorylation ^19, 20^, both of which can bind actin via both ABD1 and ABD2 domains, were destabilized by F-actin to a notably greater extent (by ∼10 °C; Fig. 3). We propose that the initial mild disengagement of ABD1-ABD2 upon ABD1 binding to F-actin (as observed in L475P_PLS2_) is followed by a stronger domain separation in PLS2_WT_ and PLS2_S5D_ due to a competitive binding of ABD2 to F-actin resulting in bundling. The L475P mutation at the actin-binding interface of ABD2 prevents its binding to F-actin, limiting the ABD1-ABD2 separation in L475P_PLS2_.

### ABD1 binds F-actin in a similar to ABD2 mode but with a smaller footprint

While the structure of ABD2 bound to F-actin is available ^37^, the attempts to solve the structure of ABD1 in complex with F-actin were unsuccessful ^55^. This is likely due to a much lower affinity of ABD1 for F-actin. To overcome these challenges, we mapped the ABD1/F-actin interface by mutagenesis to identify regions that can be used for cross-linking to F-actin using thiol-reactive reagents. Since the Cys-null construct of PLS2 does not bundle F-actin while the corresponding PLS3 construct is correctly folded and fully functional ^48^, the latter was used to generate single-Cys mutants of human plastin RD-ABD1. Human β-actin was mutated to remove the most reactive cysteine (Cys374) while introducing a cysteine in subdomain 2 (K50C) near the anticipated ABD1 binding site. The mutated actin was expressed and purified from *Pichia pastoris* ^63^. Several PLS3 constructs with single cysteines introduced at the ABD1 surface were tested for cross-linking with KC50,374CA F-actin via N,N′-1,2-phenylenedimaleimide (oPDM), N,N′-1,4-phenylenedimaleimide (pPDM), and 3,6-dioxaoctane-1,8-diyl bismethanethiosulfonate (MTS-8-MTS) homobifunctional thiol-reactive cross-linking reagents (Supplementary Fig. 4a-d). Of all the combinations of constructs and reagents, cross-linking of Q194C RD-ABD1 to K50C on actin by MTS-8-MTS provided the highest yield of the sought product (Supplementary Fig. 4a-d) and as such was selected for reconstruction by cryo-EM.

Analysis of 155,939 256-px-long segments of decorated actin filaments yielded a model of ABD1 bound to F-actin at an overall 5.1-Å resolution (Fig. 4a,b). However, the resolution for actin was higher, and the resolution of ABD1 became progressively worse the farther from the actin (Supplementary Fig. 4e). In the reconstruction, RD was not detected, pointing to its high flexibility when not stabilized by its second binding site in ABD2 ^48^. The binding interface of CH1 is oriented towards actin similarly to that of CH3 of ABD2_PLS2_ ^37^ and spectrin ^64^ (Fig. 4c). Specifically, the loop 184-189 and the first five residues of the helix 190-206 of CH1 are sufficiently close to make contacts at both sides (subdomains 1 and 3) of the hydrophobic cleft of the “i” actin subunit but also with the 44-48 segment of the D-loop (subdomain 2) of the longitudinally adjacent subunit “i+2”. Also, in subdomain 1 of the “i+2” subunit, the tip of 79-95 helix is in proximity to residues Gln118, Asn213, and His225 of plastin’s CH1. With the exception of Asn213, all these PLS3 residues are different from those determined in a previous model based on a ∼ 30-Å resolution reconstruction of ABD1-decorated actin filaments ^65^.

**Figure 4.**
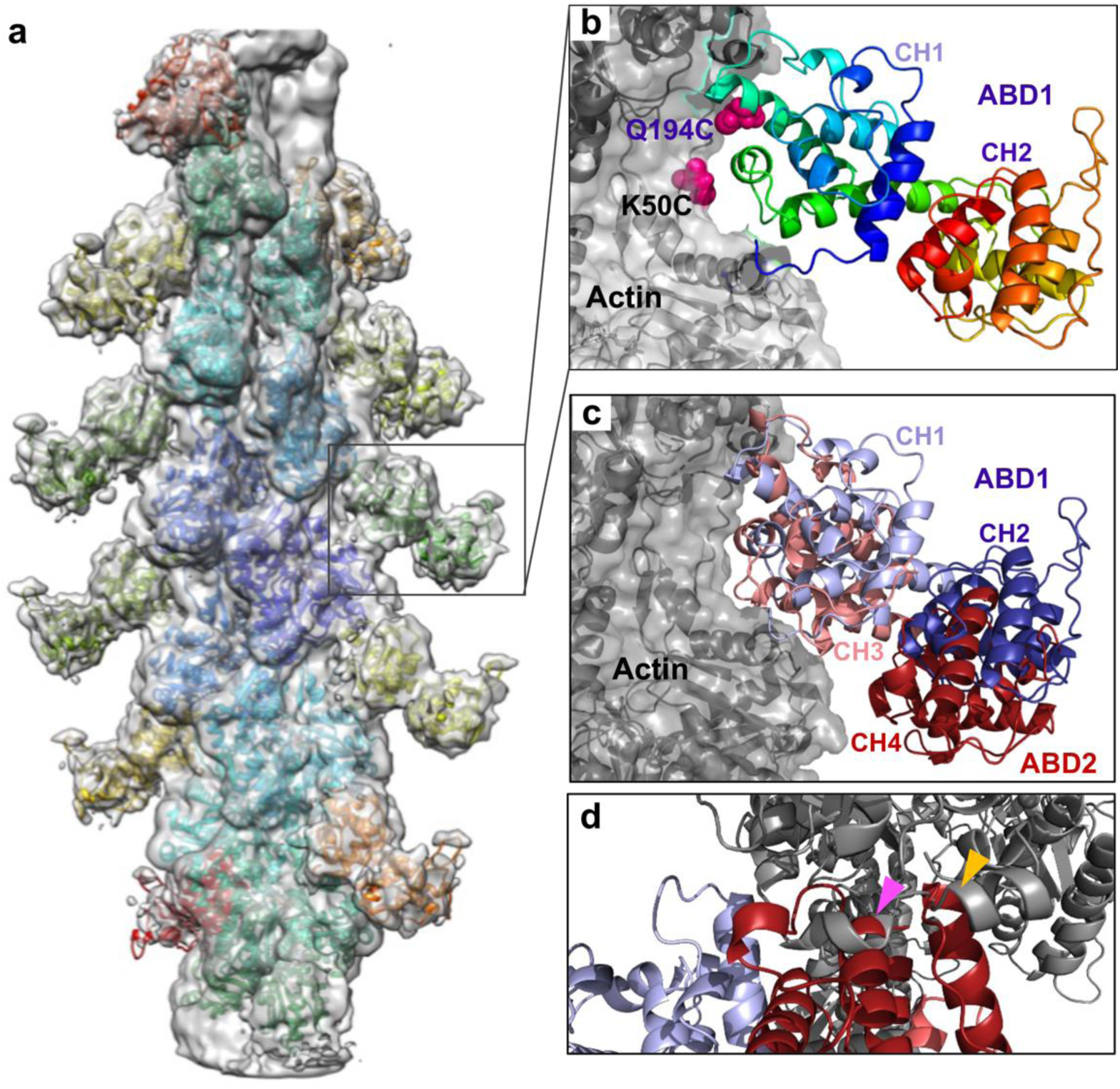
Cryo-EM reconstruction of ABD1-decorated F-actin. **(a)** Cryo-EM density map of actin filament decorated by ABD1_PLS3_ is shown as a grey transparent surface. The model built by fitting a cryo-EM structure of F-actin (PDB: 6ANU) and crystal structure of ABD1 of PLS3 (PDB: 1AOA) is shown as colored ribbons. **(b)** Close-up view of the boxed area in **a** (ABD1 is shown as rainbow-colored ribbon from N-terminus in blue to C-terminus in red; actin is in grey). Chemically cross-linked residues in ABD1 (Q194C) and actin (K50C) are shown as magenta spheres. **(c)** Superposition of ABD1 (CH1 in light blue, CH2 in dark blue; present study: EMD-25371) and ABD2 (CH3 in light red, CH4 in dark red; PDB: 6VEC) in the F-actin-bound states. **(d)** Alignment of the cryo-EM ABD1/F-actin model with the AlphaFold-generated model of human PLS3 core (https://alphafold.ebi.ac.uk/entry/P13797) revealed clashes between ABD2 and F-actin (indicated by arrowheads): 521-535 ABD2 helix (numeration as in full-length PLS2) clashes with 359-365 actin helix (magenta arrowhead); 553-562 ABD2 helix clashes with 356-358 actin loop and 359-365 actin helix (yellow arrowhead). Color scheme as in **c**.

Alignment of the cryo-EM ABD1 structure with an AlphaFold-generated model ^66^ of the PLS3 core (Fig. 4d) and the X-ray structures of yeast and plant fimbrin cores ^30^ revealed clashes between ABD2 and the actin filament. The reorientation of the ABD2 relative to ABD1 required to eliminate the clash is the likely trigger of the experimentally detected (Fig. 3) mild disengagement of ABD1 from ABD2, enabling the initial priming of ABD2 for bundling.

### A mutation at the ABD1-ABD2 interface mimicking physiologically relevant PLS2 phosphorylation reduces the relative affinity of the actin-binding domains and potentiates F-actin bundling

Since ABD1-actin affinity is relatively low (K_d_ = 2.6 µM), only a thermodynamically equivalent portion of the potent ABD1-imposed inhibition can be released upon binding to actin, bestowing a weak to moderate bundling strength. A more effective uncoupling can result in a more active protein with different properties. We sought to determine whether such modulation can be achieved by physiological means. To this end, we focused on physiologically relevant post-translational modifications of PLS2. Phosphorylation of PLS2 residues Ser5, Ser7, and Thr89 has been characterized to various degrees ^9, 13, 67, 68^, while the effects of phosphorylation at several other sites identified by high-throughput methods (PhosphoSitePlus ^69^) have not been examined. In particular, phosphorylation of Ser406 in PLS2 has been identified in several high-throughput mass-spectrometry studies in both mice and humans ^70–78^. S406 resides at the interface between the ABDs, *i.e.*, in a strategic position for a plausible regulation of ABD1/ABD2 crosstalk (Fig. 5a).

**Figure 5.**
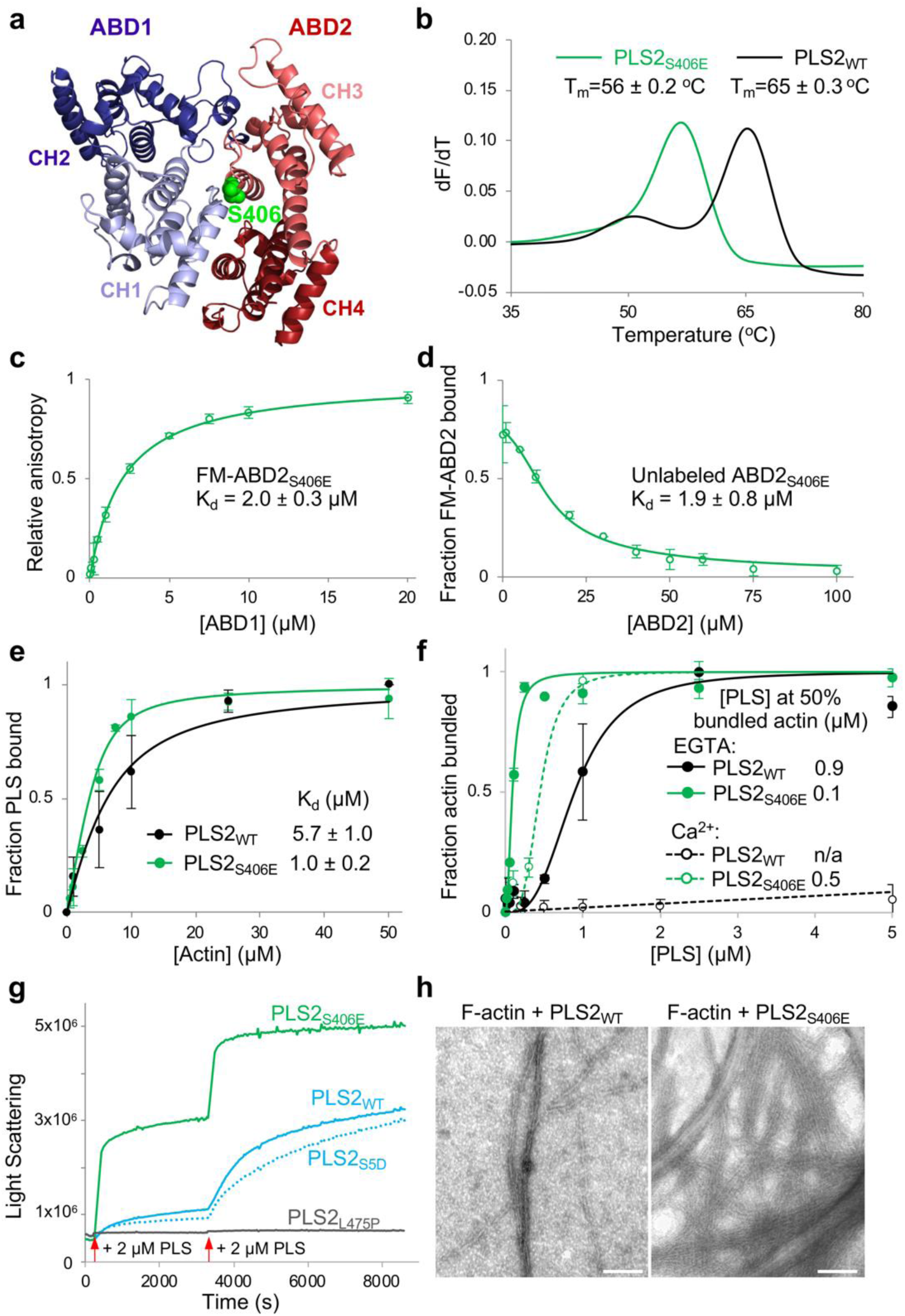
S406E PLS2 mutation mimicking physiologically relevant phosphorylation of S406 releases the inhibition of ABD2 by uncoupling it from ABD1. **(a)** An AlphaFold-generated model of PLS2-core. The location of S406E mutation is highlighted in green. **(b)** The T_m_s of WT and S406E PLS2 were assessed by DSF. The first derivative (dF/dT) traces represent the average data of n=3. **(c, d)** The binding affinity of ABD1_WT_ and ABD2_S406E_ to each other were measured by fluorescence anisotropy (**c**) and confirmed by FA competition using unlabeled ABD2_S406E_ (**d**). Error bars represent the SD of the mean; n=3. **(e, f)** The F-actin binding (**e**; n=3) and bundling (**f**; n=2) activities of PLS2_WT_ and PLS2_S406E_ were tested in high- and low-speed co-sedimentation assays, respectively. Error bars represent the SD of the mean. **(g)** F-actin bundling by WT, S5D, L475P, and S406E PLS2 was monitored by light scattering. Red arrows indicate two time points, at which the plastin constructs (2 µM final concentration) were added to F-actin. **(h)** Representative TEM micrograph images show actin bundles formed in the presence of PLS2_WT_ and PLS2_S406E_. Scale bars are 100 nm.

We produced and characterized a phospho-mimetic PLS2_S406E_ construct, which was significantly less thermostable as compared to PLS2_WT_ (T_m_ = 56 ± 0.2 °C and 65 ± 0.3 °C, respectively; Fig. 5b), tentatively reflecting a disengagement of ABD1 from the stabilizing and inhibitory interaction with ABD2. Indeed, the *in trans* affinity between the ABD1_WT_ and ABD2_S406E_ was decreased more than two orders of magnitude as compared to ABD2_WT_ (K_d_ = 2 µM vs K_d_ = 11.4 nM; Figs. 5c,d and 2b, respectively). In agreement with the inhibition hypothesis, full-length PLS2_S406E_ showed moderately improved F-actin binding and dramatically reinforced bundling efficiencies (Fig. 5e,f). Although bundling of actin by PLS2_S406E_ was inhibited by Ca^2+^, it was still stronger than bundling by PLS2_WT_ in the absence of Ca^2+^ (Fig. 5f). Therefore, Ca^2+^ binding to EF-hands cannot compensate for the phosphorylation-dependent attenuation of the allosteric cooperation between ABD1 and ABD2. The mutation strongly accelerated the rate of bundling and size of bundles as detected by light scattering and transmission electron microscopy (TEM), respectively (Fig. 5g,h). The release of the inhibition from ABD2 was not, however, complete, as under the conditions when isolated ABD2 was highly potent (*i.e.,* at 50-nM concentration (Fig. 1e)), PLS2_S406E_ showed no nucleation (Supplementary Fig. 5a), and it displayed only moderately increased nucleation activity at high (1-µM) concentration in TIRFM assays (Supplementary Fig. 5b).

Interestingly, phosphorylation analogous to that of PLS2 at S406 was not reported for either PLS1 or PLS3. In our hands, the respective recombinantly produced phospho-mimetic PLS1_S407E_ and PLS3_S409E_ constructs aggregated upon purification. Since ABD2_PLS1_ and ABD2_PLS3_ are unstable when expressed as individual domains, the observed aggregation of the phospho-mimetic mutants is likely due to the reduced stabilization of the ABD2 domains by ABD1 in these isoforms.

These data strongly support the hypothesis that ABD2 in the normal mode of plastins is potently inhibited by ABD1. This inhibition can be tuned down upon binding of ABD1 to F-actin and weakened further by physiological modifications (*e.g.,* PLS2 S406 phosphorylation). Uncoupling of the domains releases the inhibition of ABD2, resulting in its higher affinity for actin and higher bundling capacity of the protein.

### Release of ABD2 inhibition alters cellular localization of plastins

To better understand the role of the allosteric inhibition, we examined the effects of uncoupling of the ABD1/ABD2 interaction by mutations in a cellular context. Similar to WT PLS3 ^37^, WT PLS1 and PLS2 (all C-terminally tagged with mEmerald) were enriched at the cell leading edge, where they localized to actin-rich structures in the lamellipodia, filopodia, focal adhesions, and, to a lesser degree, stress fibers in *Xenopus laevis* XTC fibroblasts and human U2OS osteosarcoma cells (Fig. 6a,b and Supplementary Fig. 6). When tracked by single-molecule speckle (SiMS) TIRFM ^79, 80^, all plastin isoforms undergo retrograde flow in the lamellipodia in a similar to actin mode (rates ∼0.025 nm/s; Fig. 6c and Supplementary Video 1). In contrast, the PLS2_S406E_ construct was depleted from the lamellipodia but highly enriched at focal adhesions and stress fibers, whose morphology was enhanced and perturbed (Fig. 6a,b and Supplementary Fig. 6). Recycling of plastin from focal adhesions to the lamellipodial branched actin network requires unperturbed Ca^2+^ sensitivity ^37^. The localization of PLS2_S406E_ at the focal adhesions, therefore, is logically explained by its enhanced bundling capacities even in the presence of Ca^2+^ notably exceeding those of PLS2_WT_ under no Ca^2+^ conditions (Fig. 5f). Since PLS2_S406E_ cannot be adequately inhibited by Ca^2+^, it likely gets depleted from the lamellipodial branched actin networks via the retrograde flow leading to a concomitant accumulation at the focal adhesions and stress fibers in the lamella.

**Figure 6.**
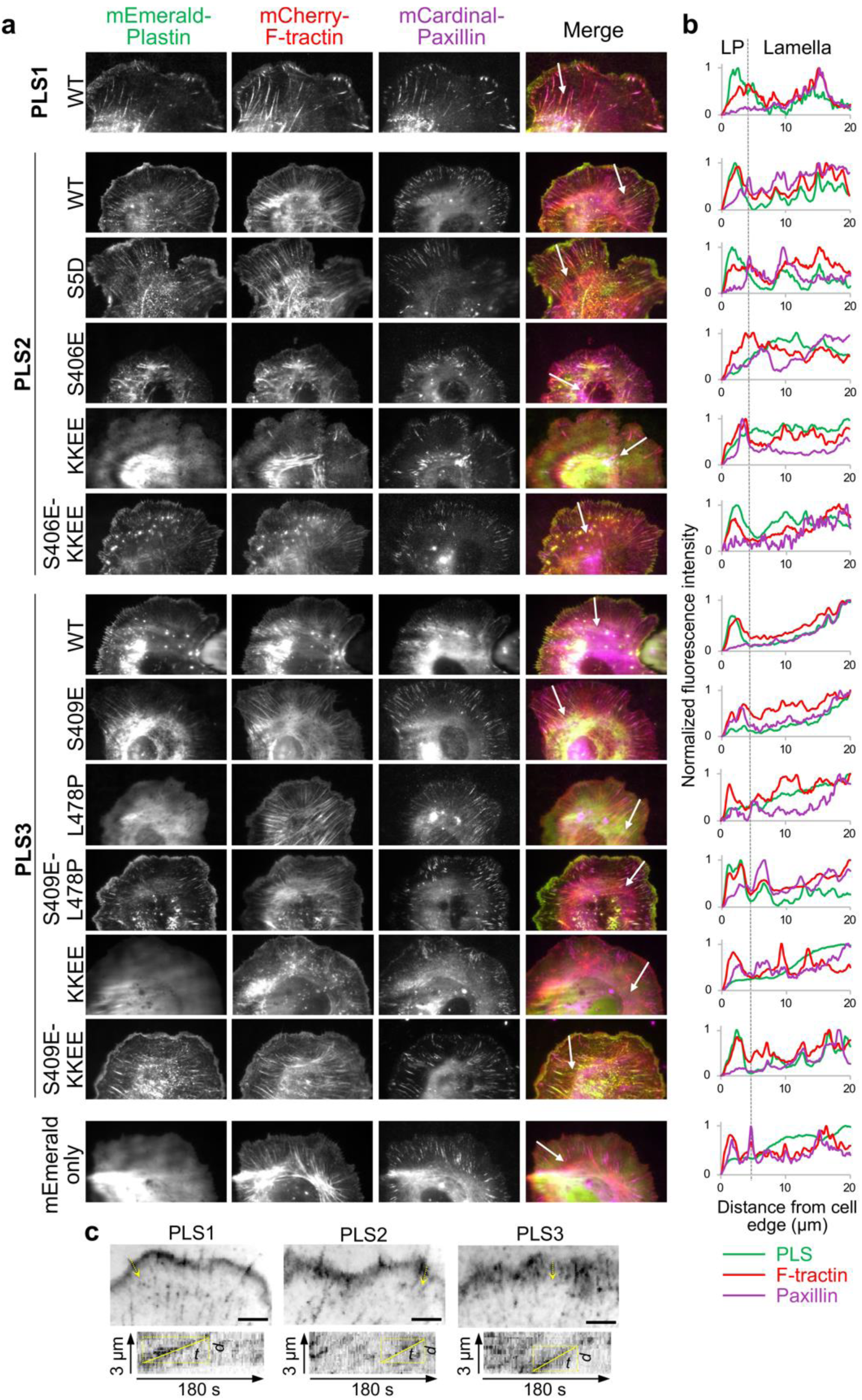
Intracellular localization of human plastins. **(a, b)** XTC cells transiently co-transfected with the indicated mEmerald-tagged plastin constructs, mCherry-F-tractin (to visualize actin) and a focal adhesion marker mCardinal-paxillin were imaged using TIRFM (**a**). Fluorescence intensity profiles (**b**) for all three channels were generated along the white arrows shown in **a**, which also serve as scale bars (20 µm). LP is lamellipodia. **(c)** SiMS TIRF microscopy of XTC cells transiently transfected with mEmerald-tagged WT plastins: average intensity projections and kymographs are shown to demonstrate retrograde flow (see also Supplementary Video 1). Velocities are calculated by dividing the distance (d) by the time (t) using bounding rectangles for the lines drawn to trace the tracks of interest on the kymographs. Scale bars are 5 µm.

Weakening the actin-binding surface of ABD2_PLS2_ by charge-inversing KK542,545EE mutations (KKEE), leading to actin-bundling deficiency *in vitro* ^48^, resulted in a diffuse localization of the mutated PLS2 and PLS3 proteins (Fig. 6a,b). Such localization is similar and consistent with that of a PLS3 congenital osteoporosis variant with the actin-binding interface of ABD2 impaired by the L478P mutation ^37^, corroborating that binding via both domains (*i.e.*, the bundling ability) is essential for plastin localization to actin-rich structures in the cell. Introduction of the ABD1-ABD2 disengaging mutation S406E on the KK542,545EE background (S406E-KKEE) restored normal localization of PLS2 to F-actin-rich structures in XTC fibroblasts (Fig. 6a,b), suggesting that a weakened ABD2 interface can be compensated by attenuating the inhibition from ABD1. In human U2OS cells, this same PLS2 construct was enriched stronger than PLS2_WT_ at focal adhesions (Supplementary Fig. 6), pointing to cell-specific and likely tissue-specific aspects of plastin physiology. Curiously, while being unstable upon expression in *E. coli*, PLS3_S409E_ localized to stress fibers in eukaryotic cells similarly to PLS2_S406E_ (Fig. 6 and Supplementary Fig. 6). This result suggests that the mutation-caused destabilization of ABD2_PLS3_ is compensated in the cell either by the presence of actin or other cellular factors.

## DISCUSSION

In a large family of CH-domain actin organizers, plastins are unique in containing two ABDs in the same polypeptide chain in intimate association with each other ^30^. Although several studies have reported that the two ABDs are not identical in their properties ^48, 55, 56, 81^, a comprehensive understanding of the domains’ mutual influence, their contribution to the function of plastins, as well as the overall logic and capacities of the molecule design have not been achieved. In this report, we addressed these deficiencies by characterizing the interaction of full-length human plastins and their individual domains with actin and with each other.

While the ability of plastins to organize aligned bundles (*e.g.,* in microvilli, cochlear stereocilia, and filopodia) is understandable and shared with other compact actin bundlers (*e.g.,* fascin), their affinity towards complex networks of poorly aligned actin filaments (*e.g.,* at the leading edge, in podosomes, immune synapses, and cuticular plates of hair cells) is puzzling. Indeed, filaments in such meshes are typically short, and the concentration of overlapping and properly oriented filaments suitable for cross-linking in such areas is low. In the present study, we discovered an allosteric inhibition between ABDs and mechanisms of its physiological attenuation resulting in dramatic changes in properties of plastins demonstrated at the molecular and cellular levels. We propose that the allosteric autoinhibition is the key feature of plastins/fimbrins underlying their versatile involvement in various cellular processes and association with both well and poorly aligned filament bundles (Fig. 7a,b).

**Figure 7.**
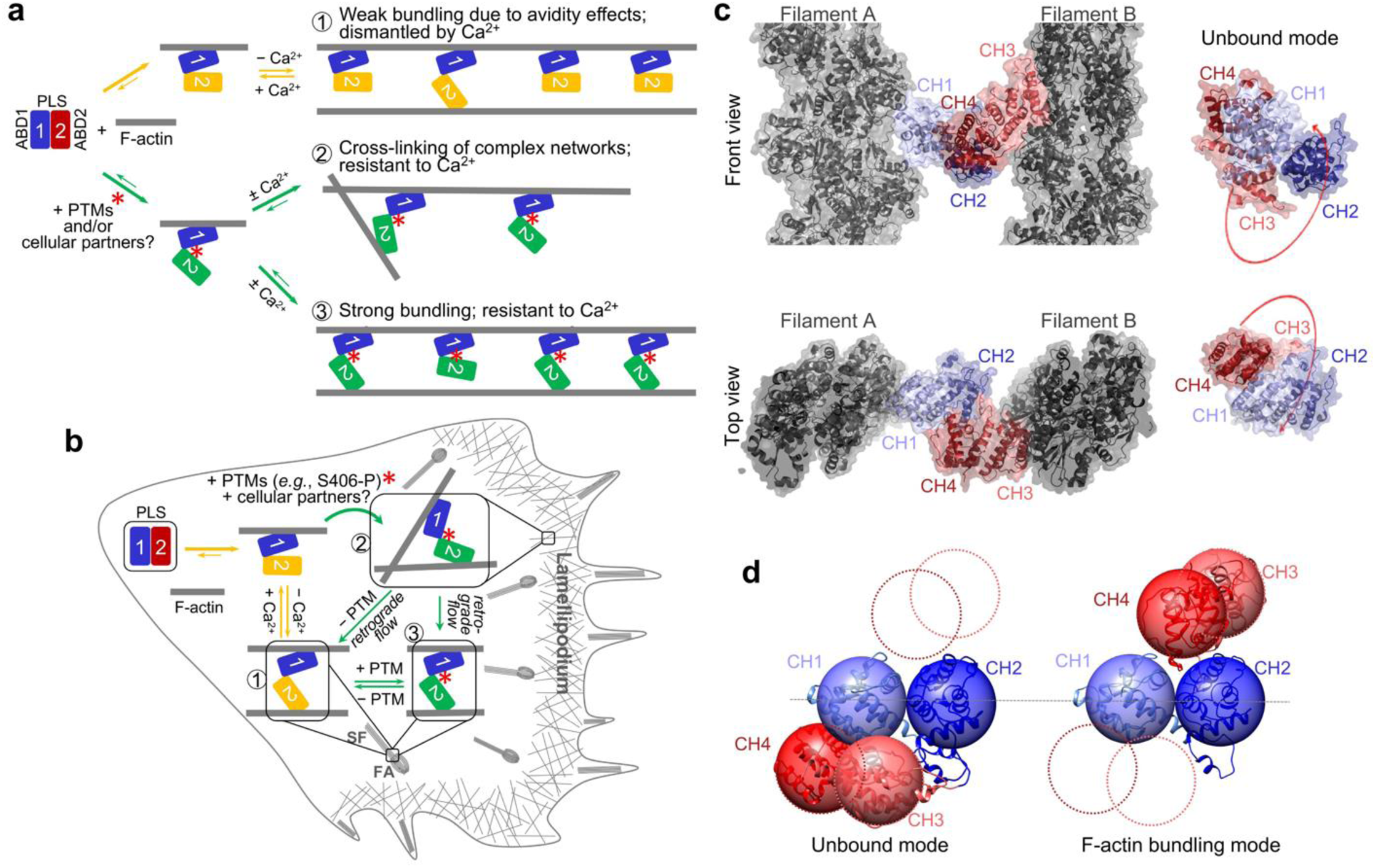
Hypothetical experimental model of actin bundling by plastins. **(a, b)** In a regular operation mode of human plastins (yellow arrows), ABD2 is strongly inhibited by ABD1. Binding of ABD1 to F-actin favors slight release of the inhibition and allows ABD2 to get engaged in weak interaction with actin (1). Due to a high affinity of ABD1 to ABD2, the inhibition is not fully released upon ABD1 binding to actin, resulting in a weak bundling mode optimal for organized bundles, where many plastin molecules are engaged in cumulative weak interactions in the F-actin bundle (1). Binding of Ca^2+^ to the regulatory domain (RD) disfavors further release of ABD2 inhibition but does not interfere with actin binding via ABD1. PTMs (*e.g.*, phosphorylation of S406) and possibly binding of unknown cellular factors weaken the ABD1-ABD2 contacts more dramatically, allowing much stronger interaction of ABD2 with actin (green arrows). This strong-bundling mode enables both, individual cross-linking events essential for strengthening branched meshes (2) and stronger, Ca^2+^-hypo-sensitive organized bundles (3). Plastins undergo retrograde flow with actin in the lamellipodia (present study), and their recycling from focal adhesions (FA) to lamellipodial branched networks is controlled by Ca^2+ 37^. In the absence of finely tuned regulation (*e.g.*, constitutively active phospho-mimicking PLS2_S406E_ and PLS3_S409E_) this eventually leads to plastin depletion from the lamellipodial branched actin networks with concomitant accumulation at the focal adhesions (FA) and stress fibers (SF) in the lamella (**b**). Under normal physiological conditions, however, delicate PTM and Ca^2+^ regulation restores the balance and allows prompt recycling of plastins from the FA/SF to the lamellipodia. **(c, d)** Model of actin bundle cross-linked by plastin based on cryo-EM reconstructions of ABD1 (EMD-25371) and ABD2 (PDB: 6VEC). The domain reorientation required for parallel, in-register actin bundle is illustrated in **d** with alternative positions of CH3-CH4 depicted by dashed circles (see also Supplementary Video 2).

All human plastins bind actin with similar affinities in a micromolar range ^48^. Despite the low affinity of full-length plastins to actin, we found that ABD2 of PLS2 alone binds actin tightly with a low-nanomolar K_d_ (Fig. 1b,c). This high affinity, however, is balanced by an equally strong inhibitory interaction with ABD1 (Fig. 2). We confirmed our prior finding ^48^ that binding of the domains to actin is sequential, and that the engagement is primed by binding of ABD1 to actin. This binding attenuates, albeit slightly, the allosteric inhibition (Fig. 3), allowing weak binding of ABD2 to an adjacent filament providing such is available in proximity. Under these conditions, ABD2 remains largely inhibited by a still very strong interaction with ABD1. We propose that this weak bundling mode is optimal for bundling of highly organized parallel actin arrays (*e.g.*, in microvilli and filopodia) where the adequate bundling strength is achieved through avidity. In this mode, plastins are highly sensitive to the inhibition and dissociation by Ca^2+^, enabling effective control of their localization ^37^ via respective signaling.

Such weak binding is not effective, however, for the organization of less ordered bundles such as those found in the branched networks associated with the lamellipodia or endocytic machinery. Additional attenuation of the allosteric auto-inhibition required in this case can be bestowed by a post-translational modification (PTM) such as PLS2 S406 phosphorylation. Since PLS2_S406E_ is a notably better bundler (Fig. 5f-h), it can cross-link poorly aligned filaments in single nodes essential for organization of meshed networks. Alternatively, attenuation can be achieved via interaction with proteins or other molecules that bind at the ABD1-ABD2 interface, *i.e.*, at a cryptic site obscured by this strong interaction. As the plastin bridges progress with retrograde actin flow towards lamella (Supplementary Video 1), their recycling to the lamellipodial leading edge is orchestrated by Ca^2+ 37^, which reaches the highest concentration at the interface between lamella and lamellipodium ^82^. Ca^2+^, however, is unable to recycle phosphorylation-mimicking PLS2_S406E_ and PLS3_S409E_ with the attenuated ABD1-ABD2 inhibition, leading to their depletion from the lamellipodia and accumulation in aligned bundles of focal adhesions and stress fibers (Fig. 6 and Supplementary Fig. 6) in agreement with a low Ca^2+^-sensitivity of PLS2_S406E_ in vitro (Fig. 5f). Therefore, our model suggests that the attenuation must be released by either removal of the PTM or dissociation from a hypothetical lamellipodial partner. In differentiated cells (*e.g.,* enterocytes and inner ear hair cells) similar mechanisms can be employed to produce stable actin bundles of microvilli and stereocilia resistant to the presence of Ca^2+^.

Interestingly, although S406 and the surrounding residues are highly conserved in all human plastins, their phosphorylation has been experimentally detected thus far only in PLS2. A cancer-centered focus of the proteomic studies detecting the S406 phosphorylation may explain this discrimination, as PLS2 was linked to carcinogenesis more than other isoforms. Alternatively, other modifications and/or interactions with protein partners may play a similar auto-inhibition attenuating role for PLS1/3. In the latter case, the localization of the partners, rather than PTM enzymes, would define the localization of plastins. Among likely candidates for such partners are coronin and mitotic spindle positioning (MISP). Indeed, coronin, which is known to be enriched at the Arp2/3 complex organized actin networks, has been reported to interact with PLS3 ^38^ and, therefore, may mediate its recruitment to lamellipodia and endocytic sites. MISP, on the other hand, has been recently shown to localize to microvilli rootlets in differentiating enterocytes and specifically recruit PLS1 to this area ^83^. The cooperativity between MISP and PLS1 facilitates cross-linking of the proximal ends of core bundles in the terminal web of brush border microvilli.

Is the extent of domain separation in plastins proposed here supported by literature data? In physiological bundles, where the role of plastin/fimbrin is unambiguously established (*e.g.,* inner ear stereocilia and intestinal microvilli), actin filaments are found in the parallel alignment. Accordingly, 2D-actin arrays assembled in the presence of plastins are also parallel ^56^. By constraining the experimentally determined distance between centers of actin filaments at 12 nm ^56, 84^ and using our cryo-EM reconstructions of actin decorated by ABD1 (Fig. 4) and ABD2 ^37^, we modeled how ABDs of plastin are expected to separate in a parallel, in-register actin bundle (Fig. 7c,d). As illustrated in Supplementary Video 2, the domain reorientation required for such assembly is rather dramatic, which, however, is in agreement with a substantial loss of thermal stability by plastins in the bundling mode (Fig. 3). Such strong separation of the domains supports a possibility for plastins to be engaged in the organization of various types of actin assemblies, including antiparallel bundles (as found in the contractile ring) and actin networks with angled filament orientations (as present in lamellipodia and endocytic patches). Experimentally, plastins/fimbrins has been found associated with both types of structures ^37, 51, 81, 85^. Furthermore, *in vitro*, at least *Saccharomyces pombe* yeast fimbrin can form actin bundles in both parallel and antiparallel orientations with a similar likelihood ^81^.

The postulated dramatic conformational changes in plastins accompanied by domain separation are consistent with recognized mechanisms of mechanosensing that operate via *i)* dynamic “catch” and “slip” bonds ^86^ and *ii)* exposure of cryptic sites in cytoskeletal proteins ^87, 88^. Indeed, several indirect but strong lines of evidence support plastin’s role in mechanotransduction. Thus, lamellipodia, endocytic patches, focal adhesions, and the contractile ring (*i.e.,* structures associated with plastins/fimbrins) are the elements whose functionality critically depends on a dynamic response to mechanical forces. Furthermore, PLS3 in mature osteocytes is highly enriched in the bifurcation sites in the osteocyte processes ^89^, *i.e.*, in regions thought to be critical for the shear-stress mediated bone remodeling. It is tempting to speculate that the anticipated sensitivity to mechanical forces underlies the important role of PLS3 as one of the main cross-linkers in microvilli and inner ear stereocilia as it may actively participate in tuning the well-recognized mechanotransduction properties of these membrane protrusions ^90^. At the molecular level, plastins have been implied in direct regulation of the transition between active and inactive forms of integrins ^9, 91, 92^, key mediators of mechanotransduction from the extracellular matrix to the cell cytoskeleton. On the other hand, it should also be noted that the proposed model of the postulated domain rearrangements is based upon separate reconstructions of individual ABD1 and ABD2 bound to actin. In these reconstructions, both CH2 and CH4 make minimal if any contacts with actin. Thus, their positions when full-length plastin is cross-linking two parallel actin filaments could be different. Thus, further experimental verification of the proposed states is required.

Since ABD2 nucleates new actin filaments (Fig. 1d,e and ^81^), the allosteric inhibition of ABD2 may also serve to prevent undesirable ABD2-induced nucleation of actin filaments. The inhibitory influence of ABD1 effectively prevents this risky outcome (Fig. 2a), and the effect sustains even when the allosteric influence is attenuated by the S406E mutation, in which case moderate nucleation can be observed only at high PLS2_S406E_ concentrations (Supplementary Fig. 5). Regardless, ABD2-imposed nucleation on the side of existing filaments is an intriguing possibility that deserves a separate investigation. Such nucleation can be used by the cells to strengthen the meshwork of branched filaments and reinforce membrane protrusions necessary to cover matrix gaps ^93^. Future studies should clarify that.

Ca^2+^ binding to EF-hands of RD is essential for plastin regulation, and altering plastin sensitivity to Ca^2+^ leads to congenital osteoporosis ^37^. Since Ca^2+^ inhibits binding of ABD2 but not ABD1 to actin ^48^, we could expect that Ca^2+^ works by strengthening the ABD1-ABD2 inhibition or by interfering with the inhibition attenuation. This hypothesis is in accordance with the predicted location of RD at the interface between ABD1 and ABD2 ^37, 48^ and with a higher affinity of RD-containing ABD1 constructs of PSL2 and PLS3 to ABD2 (Fig. 2b-e). Yet, surprisingly, Ca^2+^ showed no effect on the interactions between the actin-binding domains *in trans* (Fig. 2c,e), and only moderately affected the phosphorylation-dependent attenuation of the allosteric cooperation (Fig. 5f). These observations tentatively imply that the Ca^2+^-dependent regulation may be independent of the allosteric mechanism described here.

Given the unique properties bestowed by the tight, allosterically controlled protein design, we would expect that the unique features conferred by this distinctive domain organization are shared by all or most plastins and fimbrins. That supposition, however, is yet to be confirmed experimentally. We could not determine the affinities of ABD2 of PLS1 and PLS3 as both are not stable when expressed in isolation. However, at least in PLS3, the inhibitory influence of ABD1 on ABD2 is likely to be just as strong given the equally high affinity of ABD2_PLS2_ to ABD1_PLS3_ (Fig. 2b-e). The overall similar bundling properties of both plastins ^48^ suggest a similarly high affinity of ABD2_PLS3_ to actin. This conclusion is further supported by the similar localization of all three plastin isoforms to the leading edge of fibroblast cells (Fig. 6), and similar phenotypes of the phospho-mimicking PLS2 and PLS3 constructs (Fig. 6 and Supplementary Fig. 6).

The only other plastin family member characterized to a comparable level of detail as human PLS2 is *S. pombe* fimbrin Fim1 ^81^. Although overall Fim1 is a stronger bundler, its isolated ABD2 has much lower affinity to actin (K_d_ ∼ 650 nM) than ABD2_PLS2_ (K_d_ ∼ 4 nM; Fig. 1c). Another difference is that although ABD1 of Fim1 binds to actin similar to that of PLS2 (K_d_ ∼ 3 µM), RD-ABD1_Fim1_ binds much tighter (K_d_ ∼ 220 nM ^81^) than RD-ABD1_PLS3_ (K_d_ ∼ 4-5 µM ^48^). None of the Fim1 domains has an affinity in the low-nanomolar range. Yet, considering that ∼ 200-nM K_d_ is near the detection limit for co-sedimentation experiments, it would be interesting to reassess the affinities of the ABDs of this and other fimbrins by more sensitive approaches. If the ability to tune the cross-linking strength is an essential property of the family, we anticipate that ABD1 and ABD2 in most plastins should interact with each other with a strength comparable to that of the stronger binding domain with actin. The identity of the strongest binding domain and the sequence with which the domains interact with actin may vary.

In summary, we have characterized a novel mode of plastin regulation wherein ABD1 is always available to bind F-actin, while binding of ABD2 is secondary and a subject of regulation by the inhibitory influence of ABD1, PTMs, and Ca^2+^. We found that this regulation can be tuned by phospho-mimicking PLS2_S406E_, where S406 is a physiological PLS2 phosphorylation site. This result is of particular interest as the phosphorylation of S406 was detected in cancers ^72, 73, 75, 78^. In the context of the recognized link of PLS2 to cancer metastasis, enforced actin bundling upon S406 phosphorylation may prove to contribute to the malignant phenotype of cancer cells and thus deserves careful investigation. Future studies should clarify whether all members of the plastin family are similarly controlled by the allosteric inhibition, validate whether plastin partners (*e.g.,* coronin and MISP) can regulate plastin function and localization by tuning the engagement between ABD1 and ABD2, and test the hypothesis that exposure of cryptic sites upon bundling/domain separation is responsible for the implied mechano-sensing abilities of plastins/fimbrins.

## METHODS

### Protein expression and purification

Actin was purified from skeletal muscle acetone powder from rabbit (Pel-Freez Biologicals, Rogers, AR) or chicken (prepared in-house from flash-frozen chicken breasts (Trader Joe’s, USA)) as previously described ^94^ and stored on ice in G-buffer: 5 mM tris(hydroxymethyl)aminomethane hydrochloride (TRIS-HCl), pH 8.0, 0.2 mM CaCl_2_, 0.2 mM ATP, 5 mM β-mercaptoethanol (β-ME). Actin was used within one month with dialysis against fresh G-buffer after two weeks of storage.

All plastin constructs used in this study are described in Supplementary Table 1. To produce recombinant plastin constructs, they were cloned into pColdI vector (Takara Bio USA, Mountain View, CA) modified to include a tobacco etch virus (TEV) protease recognition site following the N-terminal 6xHis-tag ^48^. Multi-site-directed mutagenesis was carried out using the QuikChange Lightning Multi-Site-Directed Mutagenesis Kit (Agilent Technologies, Santa Clara, CA). Proteins were expressed in and purified from BL21-CodonPlus(DE3)pLysS *Escherichia coli* (Agilent Technologies, Santa Clara, CA) as described ^48^ and dialyzed against PLS buffer: 10 mM 2-[4-(2-hydroxyethyl)piperazin-1-yl]ethanesulfonic acid (HEPES), pH 7.0, 30 mM KCl, 2 mM MgCl_2_, 0.5 mM ethylene glycol-bis(β-aminoethyl ether)-N,N,N′,N′-tetraacetic acid (EGTA), 2 mM dithiothreitol (DTT), and 0.1 mM phenylmethylsulfonyl fluoride (PMSF). Purified proteins were flash-frozen in liquid nitrogen and stored at -80 °C.

### Labeling proteins with fluorescent probes

Alexa-488-actin was prepared by labeling 2 mg/ml G-actin in G-buffer devoid of reducing agents with 1.2-molar excess of Alexa Fluor 488-maleimide (Thermo Fisher Scientific, Waltham, MA) for 4 h at 4 °C, followed by dilution to 1 mg/ml and polymerization with 2 mM MgCl_2_ and 100 mM KCl overnight at 4 °C. To prepare pyrene-actin, 2 mg/ml G-actin was first polymerized in G-buffer devoid of reducing agents supplemented with 2 mM MgCl_2_ and 100 mM KCl at 25 °C for 30 min and then diluted to 1 mg/ml in F-buffer: 5 mM Tris-HCl, pH 8.0, 0.2 mM ATP, 0.2 mM CaCl_2,_ 1 mM MgCl_2_, 100 mM KCl. N-(1-pyrene)iodoacetamide (Thermo Fisher Scientific, Waltham, MA) was added to a final concentration of 40 µM, and the labeling was carried out overnight with mixing at 4 °C. Both labeling reactions were quenched with 10 mM β-ME, and labeled F-actins were pelleted at 45,000 rpm in a Ti-70 rotor in an Optima XPN 90 ultracentrifuge (Beckman Coulter, Brea, CA) for 90 min followed by three rounds of dialysis of the resulted pellets against G-buffer. All labeled and unlabeled G-actins were further purified by size-exclusion chromatography on Sephacryl S200-HR (GE Healthcare, Chicago, IL), stored on ice in G-buffer, and used within 4 weeks with dialysis to G-buffer after two weeks of storage.

ABD2_PLS2_ was labeled with fluorescein-5-maleimide (FM; Thermo Fisher Scientific, Waltham, MA). Before labeling, the protein was incubated for 1 h on ice in the presence of 10 mM DTT. The reducing agent was removed by passing twice through a NAP-5 desalting column (GE Healthcare, Chicago, IL) equilibrated with G-buffer lacking β-ME. FM was added at a 1.5-molar excess to protein and incubated overnight on ice. FM-ABD2 was passed through a NAP-5 column equilibrated with G-buffer to remove excess dye.

### Fluorescent anisotropy (FA) binding assays

G-actin was switched from Ca^2+^- to Mg^2+^-bound state by the addition of 0.1 mM MgCl_2_, and 0.5 mM EGTA and 10 min incubation on ice. Polymerization was induced by adding 2 mM MgCl_2_, 30 mM KCl, and 10 mM HEPES, pH 7.0 and incubating for 30 min at room temperature. F-actin was stabilized by a 1.2-molar excess of phalloidin (Sigma-Aldrich, St. Louis, MO). All reactions were carried out in PLS buffer supplemented with 0.2 mM ATP. For actin binding experiments, FM-ABD2 was used at 100 nM, and F-actin was added at concentrations from 0 to 2500 nM. Reactions were incubated for 30 min at 25 °C before measurement. Parallel (||) and perpendicular (|) intensities were measured on an Infinite M1000 Pro plate reader (Tecan US Inc, Morrisville, NC) with λ_ex_ = 470 nm and λ_em_ = 519 nm. Anisotropy was calculated as:

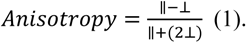

Binding affinities were determined by fitting the data to the binding isotherm equation:

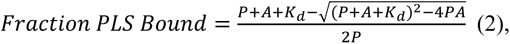

where *K_d_* is the dissociation constant, *P* is the concentration of plastin, and *A* is the concentration of F-actin.

Competition anisotropy experiments were carried out with 100 nM FM-ABD2 and 500 nM F-actin. Unlabeled ABD2 was added from 0 to 3000 nM. The dissociation constant of the unlabeled protein was determined by fitting the data to the following equation ^95^:

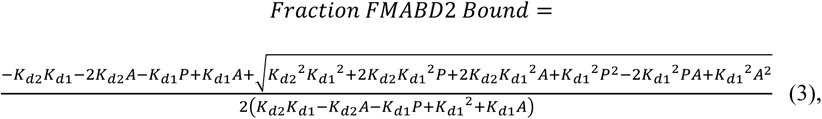

where *A* is the concentration of F-actin, *P* is the concentration of ABD2, *K_d1_* is the dissociation constant for FM-ABD2 binding to actin, and *K_d2_* is the dissociation constant for unlabeled ABD2 binding to actin. Data fitting analysis was done using Origin software (OriginLab Corporation, Northampton, MA).

Anisotropy experiments for ABD binding *in trans* were carried out similarly. The concentration of FM-ABD2 was 50 nM, and ABD1 constructs were added at concentrations from 0 to 2500 nM. In the *trans*-ABD competition assays, FM-ABD2 was at 50 nM, and ABD1 was kept at 250 nM. The concentration of unlabeled ABD2 varied from 0 to 5000 nM.

### F-actin binding and bundling co-sedimentation assays

In F-actin binding experiments carried out in PLS buffer supplemented with 0.2 mM ATP, plastin constructs were used at a final concentration of 5 μM and F-actin (polymerized as above without the addition of phalloidin) was added from 0 to 50 μM. Actin-bundling experiments contained 2 μM F-actin and plastin concentrations from 0 to 1 μM. Reactions were incubated overnight at 4 °C followed by 1 h at room temperature. Binding reactions were spun at 300,000 g for 30 min at 25 °C using Optima MAX-TL ultracentrifuge (Beckman Coulter, Brea, CA). Bundling reactions were spun at 17,000 g for 15 min at 25 °C. Supernatants and pellets were separated and analyzed by sodium dodecyl sulfate polyacrylamide gel electrophoresis (SDS-PAGE). Gels were stained with Coomassie Brilliant Blue, and band intensities were quantified by densitometry using ImageJ software ^96, 97^. Dissociation constants were determined by fitting to the binding isotherm equation (2).

Bundling efficiency was quantified by fitting the data to the Hill equation:

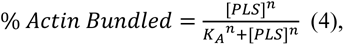

where *n* is the Hill coefficient, and *K_A_* is the concentration of plastin at 50% actin bundled.

### Bulk pyrenyl-actin polymerization assays

Ca^2+^-ATP G-actin was added to a black nonbinding 384-well plate (Corning Inc, Corning, NY) to a final concentration of 2.5 μM (5% pyrenyl-labeled). Next, F-actin was switched from Ca^2+^-bound to Mg^2+^-bound state by adding EGTA and MgCl_2_ to final concentrations of 0.5 and 1 mM, respectively. Following incubation for 1 min at room temperature, polymerization was initiated by adding 0.33 volumes of 3x initiation buffer: 30 mM MOPS, pH 7.0, 0.6 mM ATP, 1.5 mM DTT, 3 mM MgCl_2_, and 90 mM KCl. At the same time, PLS constructs were added to the desired final concentrations indicated in the figures. In inhibition experiments, ABD2 was maintained at 250 nM, and increasing concentrations of ABD1 were added. Pyrene fluorescence was monitored on an Infinite M1000 Pro plate reader (Tecan US Inc, Morrisville, NC) with λ_ex_ = 365 nm and λ_em_ = 407 nm.

### Reconstituted actin polymerization by TIRFM

For *in vitro* reconstituted TIRFM experiments, Ca^2+^-ATP G-actin (final concentration 0.9 μM, 33% Alexa-488-labeled) was switched to Mg^2+^-ATP G-actin by 2 min incubation in 0.05 mM MgCl_2_ and 0.2 mM EGTA. Actin was mixed with 50 nM or 1 µM of PLS constructs in the final reaction buffer: 10 mM imidazole, pH 7.0, 110 mM KCl, 50 mM DTT, 1 mM MgCl_2_, 1 mM EGTA, 0.2 mM ATP, 0.05 mM CaCl_2_, 15 mM glucose, 20 μg/mL catalase, 100 μg/mL glucose oxidase, 3% glycerol, and 05% methylcellulose-400cP (Sigma Aldrich, St. Louis, MO). Immediately upon mixing, reactions were transferred to a N-ethylmaleimide-myosin-treated flow chamber as described previously ^98^ and imaged using Nikon Eclipse Ti-E microscope equipped with a TIRF illumination module and a perfect focus system (Nikon Inc, Melville, NY).

### Differential scanning fluorimetry (DSF)

Heat-induced protein denaturation curves were obtained using a CFX Connect Real-Time PCR system (Bio-Rad Laboratories, Hercules, CA) as described previously ^99, 100^. Reactions were carried out in PLS buffer supplemented with 0.2 mM ATP (when actin was present) and SYPRO Orange dye (1x final concentration; Invitrogen, Carlsbad, CA). Plastin constructs were added to a final concentration of 5 μM, while actin was added at 20 μM stabilized by phalloidin at equimolar concentration. Reactions containing actin were incubated overnight at 4 °C prior to measurement. Reactions containing ABD mixtures were incubated for 30 min at 4 °C before measurement. Melting temperatures (T_m_s) were determined by calculating the max of the first derivative (-dF/dT) of the fluorescence traces (fluorescence vs temperature).

### Light scattering assays

Light scattering assays were conducted as described previously ^48^.

### Transmission electron microscopy

TEM was performed as described in ^61^.

### Sedimentation velocity analytical ultracentrifugation (SV-AUC)

SV-AUC was conducted as described previously ^101^.

### In vitro actin cross-linking by ACD toxin

ACD-catalyzed actin cross-linking was performed as described in ^61^.

### Cell culture, transfection, and microscopy

XTC cells were cultured in 70% Leibovitz’s L-15 medium (Thermo Fisher Scientific, Waltham, MA) supplemented with 10% fetal bovine serum (FBS), L-glutamine, and penicillin-streptomycin at 23 °C and ambient CO_2_. U2OS cells were cultured at 37°C with 5% CO_2_ in a humidified incubator and were grown in Dulbecco’s modified Eagle’s medium (DMEM) supplemented with 10% FBS, L-glutamine, and penicillin-streptomycin. XTC cells were obtained from Dr. Watanabe (Kyoto University Graduate School of Medicine, Kyoto, Japan) and were not further authenticated. The identity of U2OS cells was verified by STR profiling. All cells were mycoplasma-negative as determined by PCR per published protocol ^102^.

Plastin constructs (Supplemenatary Table 1) were cloned using NEBuilder (New England Biolabs, Ipswich, MA) into pcDNA3.1 with mEmerald fused at the C-termini. The plasmids pmCherry-β-actin (Addgene #54967, RRID:Addgene_54967) and pmCardinal-paxillin (Addgene #56171, RRID:Addgene_56171) were gifts from Michael Davidson ^103, 104^. Transfections were performed using Lipofectamine 3000 (Thermo Fisher Scientific, Waltham, MA). Transfected XTC cells were plated on polylysine-coated coverslips (Neuvitro Corporation, Vancouver, WA) in Attofluor chambers (Thermo Fisher Scientific, Waltham, MA) in serum-free L-15 medium and imaged 30 min post-plating. TIRFM images were obtained using a Nikon Eclipse Ti-E inverted microscope (Nikon Instruments Inc., Melville, NY, USA) equipped with a TIRF module, a perfect focus system, Nikon CFI Plan Apochromat λ 100x oil objective (NA 1.45), and an iXon Ultra 897 EMCCD camera (Andor Technology, Belfast, United Kingdom). For SiMS TIRFM, cells expressing low levels of plastins were selected, and time-lapse imaging (with 2-s intervals) was performed on small cell areas using a field diaphragm to avoid photodamage of the nucleus ^80, 105, 106^. Average intensity projections and kymographs were generated using ImageJ software. Retrograde flow of PLS constructs was estimated by kymograph analysis in ImageJ using Measure tool (bounding rectangle option): velocities were calculated by dividing the traveled distance (d) by the time (t) using the bounding rectangle for the line drawn to trace the tracks of interest on the kymographs (Fig. 6c).

U2OS cells were transfected as described above, fixed/permeabilized for 15 min in phosphate-buffered saline containing 4% formaldehyde and 0.1% Triton X-100, counter-stained with TRITC– phalloidin and Hoechst dye (Thermo Fisher Scientific), and imaged using wide-field epifluorescence, Nikon CFI Plan Apochromat λ 60x oil objective (NA 1.40), and DS-QiMc camera on Eclipse Ti-E microscope (Nikon Instruments Inc).

### Immunoblotting

MTS-8-MTS-cross-linked RD-ABD1/F-actin samples were resolved on 9% SDS-PAGE and transferred to nitrocellulose. Western blotting was performed using anti-actin (1:2000; #MA5-11869, ThermoFisher Scientific) and anti-PLS3 (1:1000; #SAB2700266, Millipore-Sigma, Burlington, MA) primary antibodies followed by anti-mouse (#A4416) and anti-rabbit (#A0545) secondary antibodies conjugated to horseradish peroxidase (both are from Millipore-Sigma and used at 1:10,000). WesternBright Sirius chemiluminescent horseradish peroxidase substrate (Advansta, San Jose, CA) was used for signal detection in Omega Lum G imaging system (Aplegen, Pleasanton, CA).

### Cryo-EM reconstruction

Cys-null PLS3 RD-ABD1 (Supplementary Table 1) was mutated to introduce a single Cys residue at positions 194 (Q194C), 216 (A216C), 226 (A226C), or 229 (A229C) using QuikChange Site-Directed Mutagenesis Kit (Agilent Technologies, Santa Clara, CA). The mutated constructs were purified as described above. *Pichia pastoris* expression vector pPICZc carrying human β-actin fused with thymosin β4 and 6xHis-tag was a gift from Mohan Balasubramanian (Addgene #111146, RRID:Addgene_111146). β-Actin cDNA was mutated to introduce K50C and C374A for labeling with a cross-linking reagent. The Cys-mutant actin was purified as previously described ^63^. In order to cross-link PLS3 RD-ABD1 to actin, each protein was reduced in the presence of 10 mM DTT for 1 h on ice before being passed twice through a NAP-5 column equilibrated with G-buffer lacking β-ME. Actin was polymerized at 3 μM as described above in the absence of DTT for 1 h at room temperature. Polymerized actin was treated with a 1.2-fold excess of a cross-linking reagent, either N,N′-1,2-phenylenedimaleimide (oPDM), N,N′-1,4-phenylenedimaleimide (pPDM), or 3,6-dioxaoctane-1,8-diyl bismethanethiosulfonate (MTS-8-MTS) (Toronto Research Chemicals, North York, ON, Canada) for 15 min before addition of 25 μM PLS3 RD-ABD1 and incubated overnight on ice. The cross-linking efficiency was verified by SDS-PAGE.

A 1.5-μL sample of cross-linked RD-ABD1/F-actin was applied to discharged lacey carbon grids and frozen with a Leica EM GP plunge freezer. Movies were collected in a Titan Krios at 300 keV equipped with a Falcon III direct electron detector, sampling at 1.4 Å/pixel. The defocus range was set to 1.5–2.5 μm, with a total dose of ∼55 electrons/Å^2^. MotionCor2 was used to motion-correct and dose-weight all the movies, followed by constant transfer function (CTF) estimation of the aligned images using the CTFFIND3 program ^107^. Images with sparse CTF estimation were eliminated. The e2helixboxer program in the EMAN2 ^108^ software package was used for boxing 256-px-long filaments (total 155,939 boxed segments). Relion was used for the following helical reconstruction procedure. The overall resolution of the final reconstruction was determined by the Fourier shell correlation (FSC) between two independent half maps, which was 5.1 Å at FSC = 0.143. The model of F-actin (PDB: 6ANU) and ABD1 of PLS3 (PDB: 1AOA) were fit into the cryo-EM map, followed by examination and adjustment in COOT ^109^. The manually curated model was then real-space refined in PHENIX ^110^. The cryo-EM map was deposited with accession code EMD-25371 in the Electron Microscopy Data Bank (EMDB).

### Statistical analysis

All graph data is presented as mean ± standard deviation (SD) or standard error (SE) as indicated in the figure legends. Number of repetitions (n) is indicated in the figures/figure legends. ANOVA followed by multiple comparison tests (Student’s t-tests with a two-tailed distribution) with Bonferroni correction was applied to determine statistically significant differences; individual *p*-values are indicated in the figure legends, where appropriate.

## ACKNOWLEDGMENTS

We thank Dr. Fenbin (Jerry) Wang (University of Virginia) for the assistance in modeling parallel actin filaments linked by a plastin bridge. This work was supported by the National Institute of General Medical Sciences of the NIH under award numbers R01GM114666 (to DSK) and R35GM122510 (to EHE) and a 2018 Pelotonia Graduate Fellowship Award at The Ohio State University Comprehensive Cancer Center (to CLS). The content is solely the responsibility of the authors and does not necessarily represent the official views of the National Institutes of Health.

## AUTHOR CONTRIBUTIONS

Conceptualization – DSK; Funding acquisition – DSK, EHE, CLS; Supervision – DSK, EHE; Investigation – CLS, EK, RA, WZ; Formal analysis – CLS, EK, WZ; Visualization – EK, CLS, WZ; Writing – original draft CLS, DSK, EK; Writing – review & editing – all authors.

## DATA AVAILABILITY

The cryo-EM map of ABD1_PLS3_/F-actin complex was deposited with accession code EMD-25371 in the EMDB.

