## Supplemental information for "Allosteric Regulation of Human Plastins"

### Supplementary Figure 1

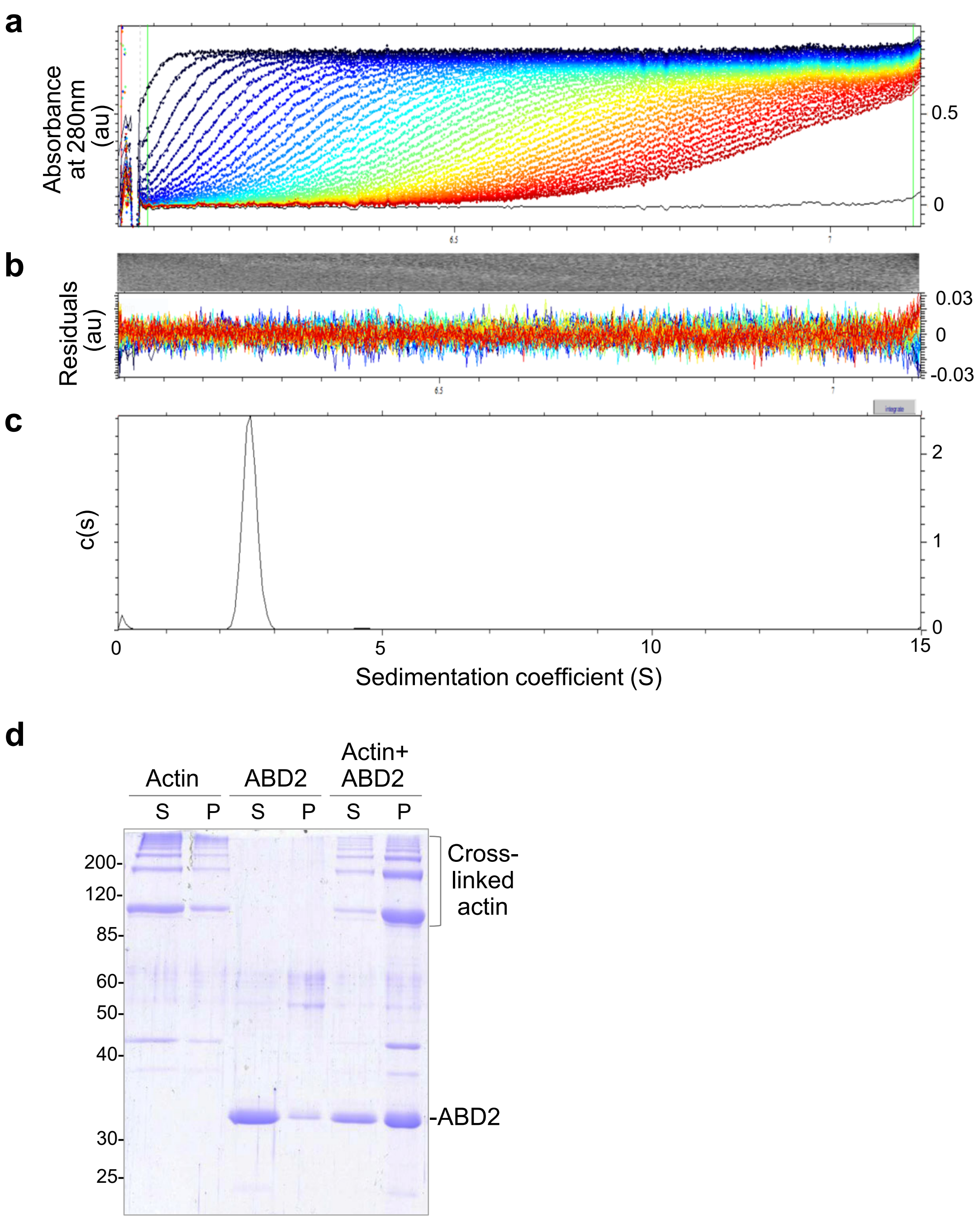


**Supplementary Figure 1 (related to Figure 1). ABD2$_{PLS2}$ exists in solution as a monomer and rescues polymerization of ACD-crosslinked actin oligomers**

**(a-c)** Sedimentation velocity analytical ultracentrifugation (SV-AUC) analysis of ABD2$_{PLS2}$. Raw sedimentation profiles of absorbance at 280 nm versus radius (**a**) and residual plots (**b**) are shown. The distribution of sedimentation coefficients (**c**) indicates the presence of only monomeric species of ABD2$_{PLS2}$ protein.

**(d)** ABD2$_{PLS2}$ rescues polymerization of ACD-cross-linked actin oligomers. G-actin was covalently cross-linked by addition of ACD toxin in the absence (Actin) or presence (Actin +ABD2) of ABD2. Following ACD treatment, cross-linked actin was allowed to polymerize by addition of $Mg^{2+}$ and KCl and subjected to ultracentrifugation to separate non-polymerized soluble (S) and polymerized pellet (P) fractions on SDS-PAGE. ABD2 alone sample (ABD2) treated identically served as a negative control.

### Supplementary Figure 2

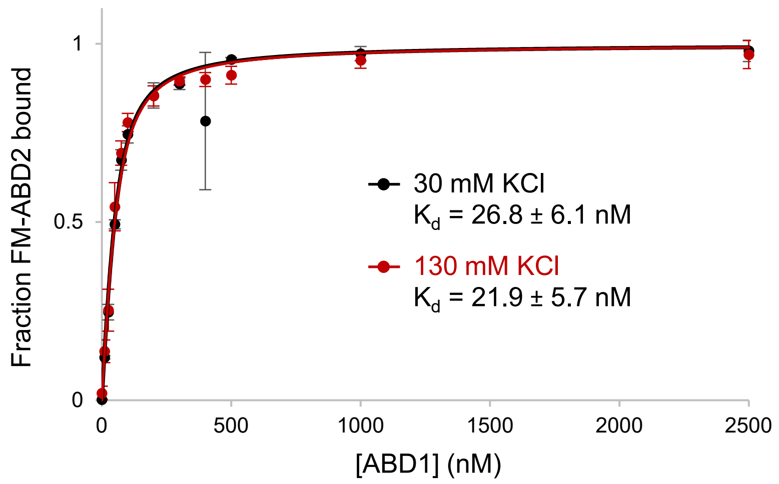


**Supplementary Figure 2 (related to Figure 2). ABD1/ABD2 interaction is not affected by salt**

The ABD1-ABD2 affinity in the presence of 30 and 130 mM KCl was determined by fluorescence anisotropy assays. Error bars represent the SD of the mean; n=3.

### Supplementary Figure 3

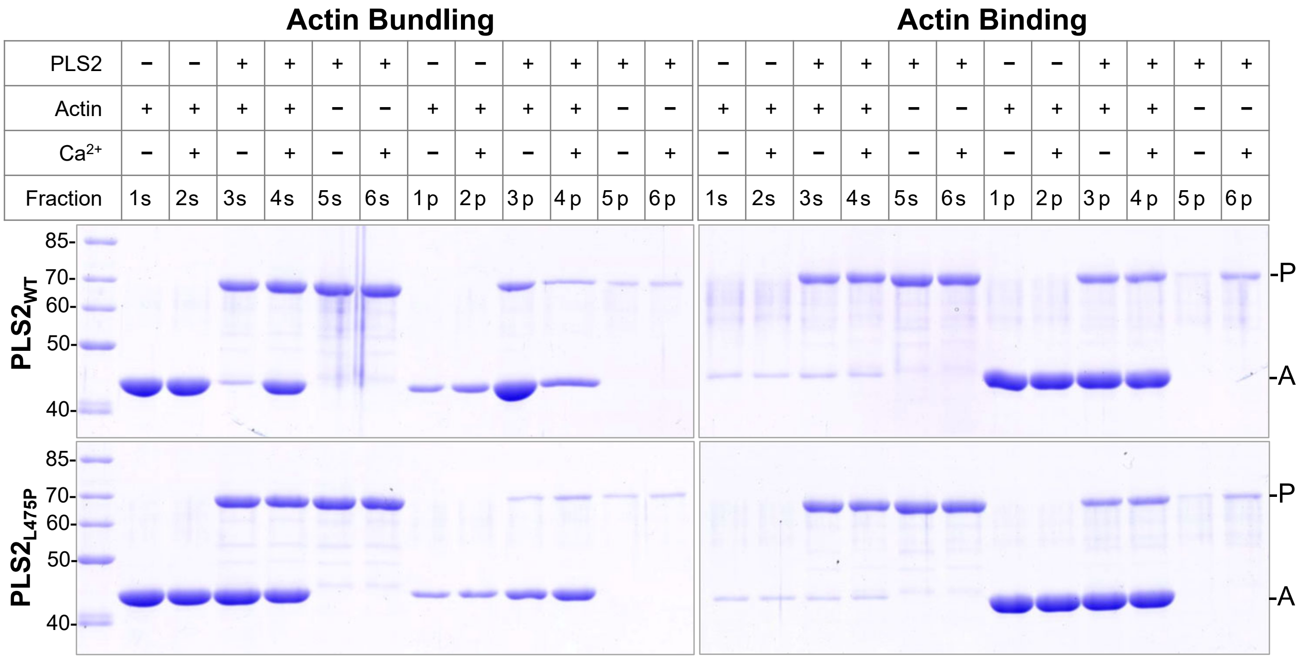


**Supplementary Figure 3 (related to Figure 3). L475P mutation diminishes F-actin bundling ability of PLS2**

F-actin bundling and binding by PLS2$_{WT}$ and PLS2$_{L475P}$ was assessed by low- (bundling) and high- (binding) speed sedimentation. Low-speed co-sedimentation (panel **Actin Bundling**) detected F-actin bundles formed only in the presence of PLS2$_{WT}$ in the absence of Ca$^{2+}$ (PLS2$_{WT}$ Actin Bundling **3s/3p** vs **4s/4p**), while PLS2$_{L475P}$ was unable to bundle F-actin regardless of Ca$^{2+}$ [F-actin remained mainly in the supernatant (PLS2$_{L475P}$ Actin Bundling **3s/3p** and **4s/4p**)]. In high-speed co-sedimentation assays (panel **Actin Binding**), both constructs were co-pelleted with F-actin in the absence (Actin Binding **3s/3p**) and presence of Ca$^{2+}$ (Actin Binding **4s/4p**) implying that L475P mutation does not affect F-actin binding mediated through ABD1.

### Supplementary Figure 4

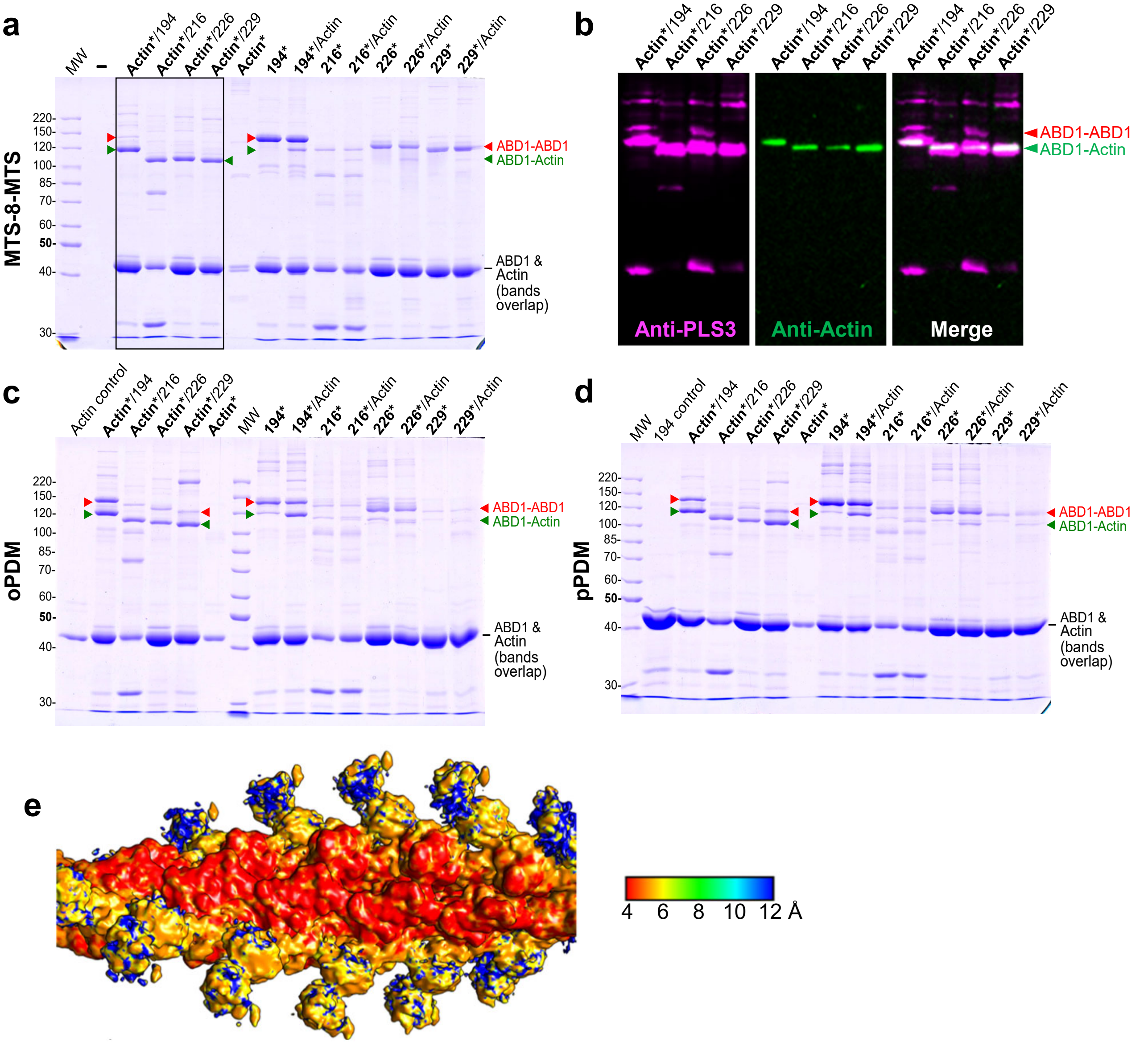


**Supplementary Figure 4 (related to Figure 4). Cryo-EM reconstruction of ABD1/F-actin**

**(a-d)** Preparation of ABD1/F-actin sample for cryo-EM. To produce F-actin decorated with ABD1, recombinant human β-Actin carrying K50C and C374A mutations was purified from *Pichia pastoris* and polymerized as described in online Methods. Individual Cys residues were introduced on the Cys-null RD-$ABD1_{PLS3}$ background (Supplementary Table 1) at the indicated positions resulting in constructs containing single cysteines (**Q194C**, **A216C**, **L226C**, or **G229C**). Asterisks indicate activation of either actin (**Actin***) or RD-ABD1 constructs (**194***, **216***, **226***, or **229***) using a corresponding cross-linking reagent [**MTS-8-MTS** (**a**), **oPDM** (**c**), or **pPDM** (**d**)]. Following the activation, 2.5 µM actin was mixed with 10-molar excess of an RD-ABD1 construct. The resulting cross-linked samples were resolved on 9% SDS-PAGE (**a**, **c**, **d**). Note that non-crosslinked actin (42 kDa) and RD-ABD1 constructs (43 kDa) have similar molecular weights resulting in similar mobility on SDS-PAGE. Anti-actin and anti-PLS3 western blotting was performed to confirm identity of the resulting bands; only the immunoblots for the samples boxed in **a** are shown in **b**. Red arrowheads indicate RD-ABD1/RD-ABD1 cross-links. Green arrowheads indicate successful formation of RD-ABD1/F-actin cross-links. The cross-linking of MTS-8-MTS-activated actin with the RD-ABD1 (boxed in **a**) was the most efficient as compared to other tested reagents/combinations, and the sample of MTS-8-MTS-activated actin cross-linked with RD-$ABD1_{Q194C}$ was subjected to cryo-EM.

**(e)** ABD1/F-actin density map is colored according to the resolution in angstroms.

Supplementary Figure 5

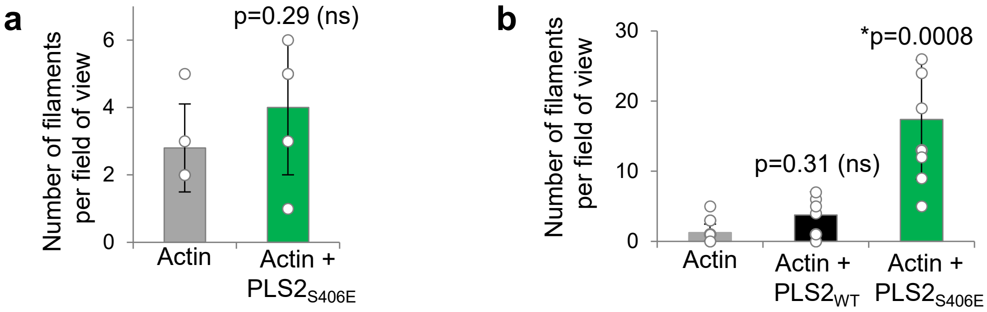


**Supplementary Figure 5 (related to Figure 5). S406E phospho-mimetic mutation moderately increases nucleation activity of PLS2**

Actin nucleation activity of $PLS2_{S406E}$ at 50-nM (**a**) and 1-μM (**b**) was tested by TIRFM. Error bars represent the SD of the mean calculated from two independent experiments each containing 4 replicates. ANOVA followed by multiple comparison tests with Bonferroni correction was applied: asterisk indicates statistically significant difference (*p<0.025) compared to the actin control.

Supplementary Figure 6

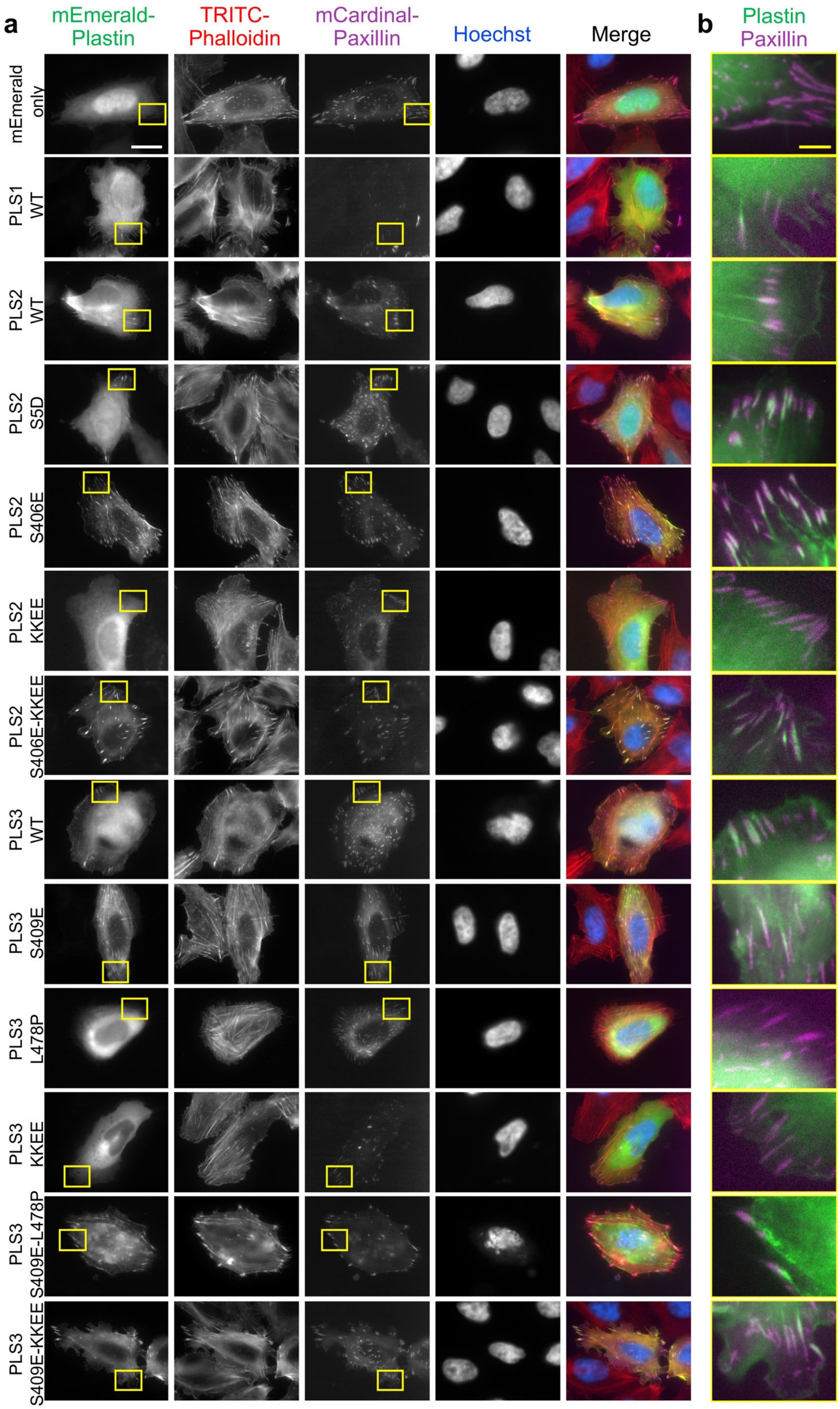


**Supplementary Figure 6 (related to Figure 6). Intracellular localization of human plastins**

U2OS cells transiently co-transfected with the indicated mEmerald-tagged plastin constructs and a focal adhesion marker mCardinal-paxillin were fixed and counter-stained with TRITC-phalloidin and Hoechst. Boxed areas in (**a**) are enlarged in (**b**). Scale bars are 20 µm in (**a**) and 5 µm in (**b**).

### Supplementary Table 1

| Construct | Residues | Mutations | Vector | Figures | |
|---|---|---|---|---|---|
| | | | | Main | Suppl. |
| **PLS1** | | | | | |
| $PLS1_{WT}$-mEmerald | 1-629 | N/A | pcDNA3.1 | 6 | 6 |
| **PLS2** | | | | | |
| $FL_{PLS2}/PLS2_{WT}$ | 1-627 | N/A | pColdTEV | 1,3,5 | 3,5 |
| $ABD1_{PLS2}$ | 87-379 | N/A | pColdTEV | 1,2,5 | 2 |
| $ABD2_{PLS2}$ | 380-627 | N/A | pColdTEV | 1,2 | 1,2 |
| $RD\text{-}ABD1_{PLS2}$ | 1-379 | N/A | pColdTEV | 2 | |
| $PLS2_{S5D}$ | 1-627 | S5D | pColdTEV | 3,5 | |
| $PLS2_{L475P}$ | 1-627 | L475P | pColdTEV | 3,5 | 3 |
| $PLS2_{S406E}$ | 1-627 | S406E | pColdTEV | 5 | 5 |
| $ABD2_{S406E}$ | 380-627 | S406E | pColdTEV | 5 | |
| $PLS2_{WT}$-mEmerald | 1-627 | N/A | pcDNA3.1 | 6 | 6 |
| $PLS2_{S5D}$-mEmerald | 1-627 | S5D | pcDNA3.1 | 6 | 6 |
| $PLS2_{S406E}$-mEmerald | 1-627 | S406E | pcDNA3.1 | 6 | 6 |
| $PLS2_{KKEE}$-mEmerald | 1-627 | KK542,545EE | pcDNA3.1 | 6 | 6 |
| $PLS2_{S406E\text{-}KKEE}$-mEmerald | 1-627 | SKK406,542,545EEE | pcDNA3.1 | 6 | 6 |
| **PLS3** | | | | | |
| $ABD1_{PLS3}$ | 92-389 | N/A | pColdTEV | 2 | |
| $RD\text{-}ABD1_{PLS3}$ | 1-389 | N/A | pColdTEV | 2 | |
| $RD\text{-}ABD1_{Q194C}$ | 1-389 | CysNull* + Q194C | pColdTEV | 4 | 4 |
| $RD\text{-}ABD1_{A216C}$ | 1-389 | CysNull* + A216C | pColdTEV | | 4 |
| $RD\text{-}ABD1_{L226C}$ | 1-389 | CysNull* + L226C | pColdTEV | | 4 |
| $RD\text{-}ABD1_{G229C}$ | 1-389 | CysNull* + G229C | pColdTEV | | 4 |
| $PLS3_{WT}$-mEmerald | 1-630 | N/A | pcDNA3.1 | 6 | 6 |
| $PLS3_{S409E}$-mEmerald | 1-630 | S409E | pcDNA3.1 | 6 | 6 |
| $PLS3_{L478P}$-mEmerald | 1-630 | L478P | pcDNA3.1 | 6 | 6 |
| $PLS3_{S409E\text{-}L478P}$-mEmerald | 1-630 | SL409,478EP | pcDNA3.1 | 6 | 6 |
| $PLS3_{KKEE}$-mEmerald | 1-630 | KK543,545EE | pcDNA3.1 | 6 | 6 |
| $PLS3_{S409E\text{-}KKEE}$-mEmerald | 1-630 | SKK409,543,545EEE | pcDNA3.1 | 6 | 6 |

**Supplementary Table 1. PLS constructs used in the present study**
N/A – not applicable.
* In CysNull PLS3 construct the following nine Cys residues were substituted by Ala: 33, 104, 143, 167, 209, 349, 463, 566, 621.

**Supplementary Videos**

**Supplementary Video 1 (related to Figure 6c). Human plastin isoforms undergo retrograde flow in the lamellipodia**
SiMS TIRFM time-lapse imaging of XTC cells transiently transfected with mEmerald-tagged human plastin isoforms. Scale bars are 5 μm.

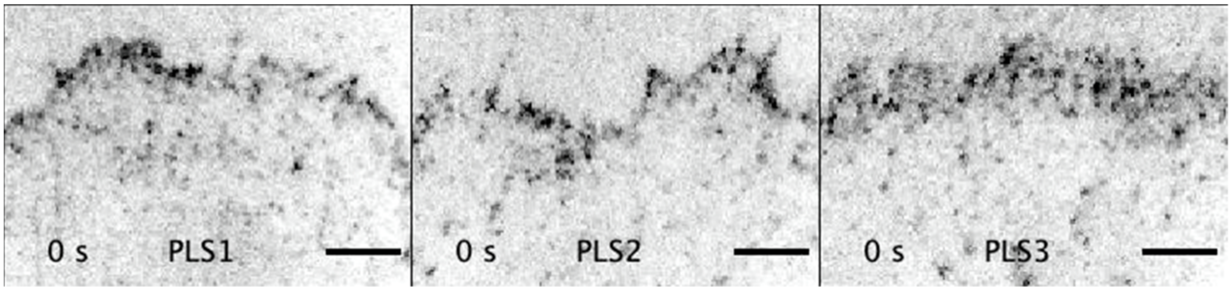


**Supplementary Video 2 (related to Figure 7d). The domain reorientation is required for parallel, in-register actin bundle**
Two modes of plastin are shown: the AlphaFold model of plastin core in the unbound state and the model for a crossbridge in actin bundle. CH1 is light blue, CH2 is dark blue, CH3 is light red, CH4 is dark red, actin is grey.

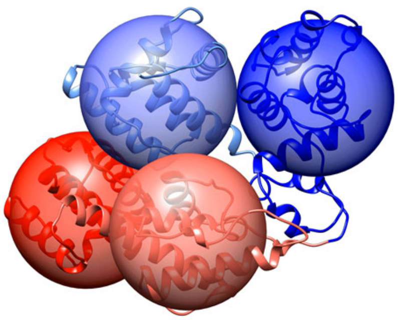
